# Cell type diversification and phenotype convergence underlying white fin-ornamentation of cyprinid fishes

**DOI:** 10.64898/2025.12.26.696501

**Authors:** Delai Huang, Tiffany Liu, August A. Carr, Pietro H. de Mello, Yipeng Liang, Leah P. Shriver, François Chauvigné, Stephen L. Johnson, Joan Cerdà, Gary J. Patti, David M. Parichy

**Author notes:** Deceased (Stephen L. Johnson).

## Abstract

Neural crest derived cells offer valuable opportunities to dissect mechanisms of cell fate specification and differentiation within individual ontogenies and the underpinnings of cell type diversification over evolutionary time. Particularly useful for such analyses are pigment cells of ectothermic vertebrates that arise from neural crest cells, or via latent neural crest derived stem cells, and comprise several classes with cell-type specific pigmentary phenotypes. Among these are white cells, “leucophores,” present in a variety of species that contribute to patterns on the body or ornamentation on the fins. To better understand developmental and evolutionary origins of these cells we have examined leucophores harboring deposits of yellow/orange carotenoids, “xantholeucophores,” of zebrafish and leucophores of white cloud minnow, within the same family Cyprinidae. We show that white phenotypes of both cell types require sepiapterin reductase that promotes accumulation of pale and colorless pteridines. We further demonstrate that xantholeucophores develop directly from yellow sepiapterin rich xanthophore-like cells and that this transition requires both gap junctional activity and permeability of the aquaglyceroporin / peroxiporin channel Aquaporin 3. These findings identify these white cells as distinct developmentally, genetically, and biochemically from another type of white cell in zebrafish and other white cells present across phylogenetic lineages. Our results highlight remarkable convergences and parallelisms in the acquisition of white cell phenotypes within and between species and identify this as a rich system for enquiries into the evolutionary individuation of novel cell types.

**Significance:** Understanding how cell types arise is fundamental to explaining animal complexity. Pigment cells offer opportunities to address this question because they display striking variation. We show that white pigment cells comprise multiple classes with independent origins. In zebrafish, white cells on one fin develop from melanophores whereas white cells on another fin develop from yellow precursors that convert their pigments to colorless ones. White cells of a related minnow require the same colorless pigments yet differ in other ways. These findings illuminate remarkable convergence: white cells have arisen repeatedly from different progenitor cell lineages by different mechanisms. This diversity—even within and between closely related species—identifies a powerful model for understanding the evolution of cell types with distinct identities and functions.

## Introduction

Organisms evolve as integrated units yet there is a renewed appreciation for cell types as evolutionary units in their own right (1–3). Elucidating how new cell types arise—with unique functional, morphological, and molecular characteristics, and distinct gene regulatory connections—is foundational to understanding organismal complexity, some adaptations, and likely patterns of cladogenesis. Recent efforts have provided conceptual advances and empirical insights into cell type evolution, emphasizing gene regulatory modules that allow individuation of new cell types from progenitors that are ancestral evolutionarily and antecedent developmentally. In some cases, molecular mechanisms leading to diversification within a cell lineage have been described; in others, convergence of phenotypes from distinct cell lineages (4–7). Yet an important question arises: how many distinct types of cells comprise even well-studied cell lineages and to what extent are they conserved phylogenetically? Such cellular taxonomy, within and between organisms, is foundational to dissecting cell type evolution itself (8, 9).

Especially amenable to studying cell type diversification over developmental and evolutionary time are neural crest derived cells of vertebrates that generate neurons and glia of the peripheral nervous system, bone and cartilage of the craniofacial skeleton, and many other cell types that contribute to many other traits (10). Neural crest derived cells are also interesting for acquiring characteristics of other cell lineages entirely, providing examples of evolutionary plasticity in cell type specification and convergence in cell function even across germ layers.

In these contexts of cell type evolution, functional diversification, and phenotypic convergence, particular opportunities are afforded by pigment cells that arise directly from embryonic neural crest cells, or indirectly via latent stem cells of neural crest origin (11–13). Pigment cells of mammals and birds are limited to melanocytes. Yet, pigment cells of ectothermic vertebrates comprise several types including melanophores, yellow/orange xanthophores, red erythrophores and iridescent iridophores (14, 15). Collectively referred to as “chromatophores,” their patterns on the body and in extremities such as fins have roles in intraspecific and interspecific signaling, camouflage, thermoregulation and other activities (16). This adaptive significance, and the tendency for pigment cells to be relatively uncoupled from pathways essential for early embryogenesis or subsequent viability (17), provides fertile ground in which to uncover new cell type diversity.

Given these considerations we have examined white cells of zebrafish and other teleosts. Such cells occur in disparate species and contribute to body patterns and fin ornamentation (14). Distinctive pattern elements on fins often serve as visual signals in courtship, dominance displays and other interactions (18–20); similarly contrasting ornaments occur in species ranging from damselflies to lizards and often have clear ethological significance (21, 22). In zebrafish, white pigment cells at the edge of the dorsal fin contribute to shoaling behavior and are especially prominent during agonistic lateral displays, in which fins are flared and presented to rivals (23, 24). These white cells—melanoleucophores—develop from melanophores that lose their melanin and acquire irregularly shaped and arranged crystals of guanine that confer a matte white phenotype (23, 25).

In this study, we focused on another fin pigment cell of zebrafish, found in reiterative light stripes of the anal fin. We show that these cells arise from yellow/orange xanthophore-like progenitors, and that transition to a “xantholeucophore” phenotype requires gap junctional communication and permeability of a transmembrane channel for water balance, glycerol and peroxide. By transcriptomic, chemical and mutational analyses, we demonstrate that the white phenotype depends on processing of yellow pteridines to pale and colorless pteridines, mediated in part by sepiapterin reductase. Finally, we show that white cells of another minnow also harbor pale and colorless pteridines requiring sepiapterin reductase activity, despite these cells having other features distinct from xantholeucophores, and occurring in an anatomical location like that of zebrafish melanoleucophores. Our study reveals a striking diversity in white cell phenotypes, across sublineages of neural crest cells and phylogenetic lineages of teleosts.

## Results

### White Fin-Pattern Features and Cells of Zebrafish and White Cloud Minnow

The zebrafish, *Danio rerio*, has prominent white ornamentation along its dorsal fin edge and at the tips of its caudal fin because white melanoleucophores (ML) differentiate from melanophores (M) at these locations (Fig. 1*A*, upper) (23, 25). “Off-white” pattern features are evident in the anal fin, too, where xantholeucophores (XL) occur in light “interstripes” that alternate with dark stripes of melanophores and unpigmented (“cryptic”) xanthophores (Xc) (26, 27). XL contain a white material and yellow/orange carotenoids and contribute a yellowish-white color to the pattern (23). Interstripes of the fin contain mostly XL with a few iridophores (I) and so differ from interstripes of the body that consist of yellow/orange xanthophores overlying very densely packed iridophores (28).

**Fig. 1.**
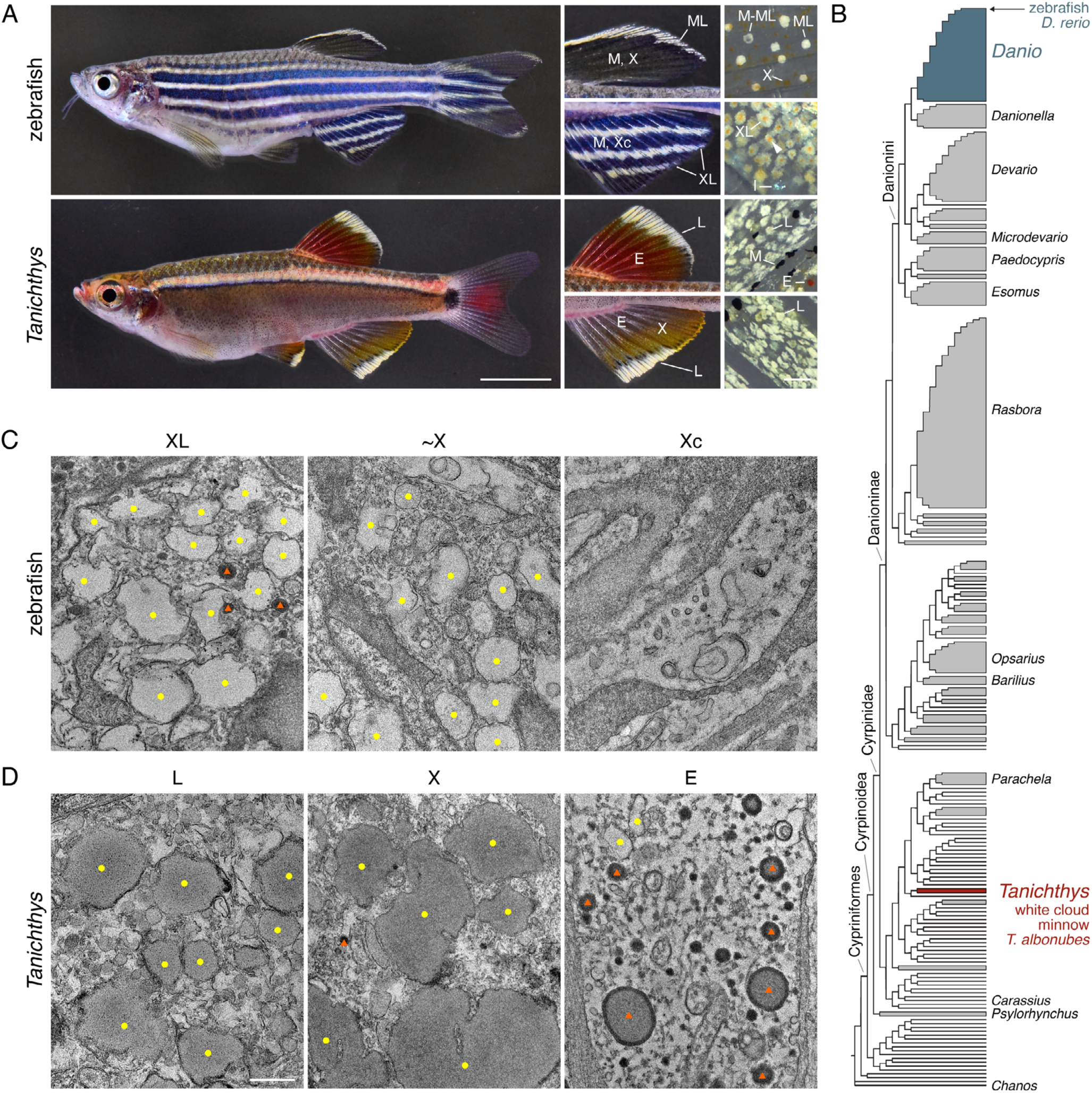
White cell types and morphologies. (*A*) Phenotypes of zebrafish *Danio rerio* (upper) and white cloud minnow *Tanichthys albonubes* (lower). For each species, middle panels show higher magnification views of dorsal fin (upper) and anal fin (lower) and far right panels show details of chromatophore pigments after treating fish with epinephrine to contract pigment granules towards cell centers). In zebrafish, melanoleucophores (ML) contain crystalline guanine that contributes white ornamentation to dorsal and caudal fins tips. These cells develop from melanophores (M), and transitional cells (M-ML) still bear some melanin in dark central spots (close-up at far right). Though confined to these locations in zebrafish, other *Danio* species also have ML in the anal fin (23). Xantholeucophores (XL) occur in light “interstripes” of the anal fin, between dark stripes of melanophores (M) and cryptic xanthophores (Xc). XL contain white material as well as yellow/orange carotenoids (23) evident as centrally located accumulations of brightly colored vesicles (arrowhead, far right; I, iridophore). In white cloud minnow, *T. albonubes*, white leucophores (L) resembled ML in occurring at the edges of the dorsal fin and anal fin and in lacking spots of carotenoids. Yet these cells also lacked persisting melanin typical of M–ML transition. (*B*) Phylogenetic relationships, showing *Danio* and *Tanichthys* within family Cyprinidae (33). (*C*) In the juvenile zebrafish anal fin (14 mm standardized standard length; 14 SSL)(69), organelles resembling pterinosomes (yellow dots) were abundant in newly differentiating XL and yellow xanthophore-like cells (∼X) present at this and earlier stages (see below); cryptic xanthophores (Xc) within fin stripes had pigmentary organelles only at early stages of biogenesis. Structures marked with red triangles are likely to be lipid droplets, and potentially carotenoid-containing vesicles. (*D)* In anal fins of adult *Tanichthys*, pterinosome-like organelles were apparent in leucophores (L) and xanthophores (X), and presumptive carotenoid vesicles and pterinosomes were present in erythrophores (E) (54). (Scale bars: *A* 5 mm and 50 µm, *D* for *C*, *D* 500 nm).

ML and XL occur across *Danio* species (23, 25) whereas a different type of leucophore occurs in medaka, *Oryzias latipes* (29, 30). *Danio* and *Oryzias* belong to different superorders that diverged 213–280 million years ago (31, 32). We asked whether these or other types of leucophores might occur in species more closely related to zebrafish, but outside of *Danio*. We therefore examined white cloud minnow, *Tanichthys albonubes*, hereafter *Tanichthys*, a member of the same family as zebrafish, Cyprinidae, but in a lineage that diverged from that of zebrafish 32–52 million years ago (Fig. 1*B*) (33, 34).

*Tanichthys* dorsal and anal fins have white patches along their edges with a similar appearance to that of the zebrafish dorsal fin, where ML reside (Fig 1*A*, lower). Yet *Tanichthys* white cells—leucophores—within these patches lacked discernible melanin, as in transitional or differentiation-arrested ML. These cells also lacked spots of yellow/orange carotenoids, as seen in XL following epinephrine treatment. Instead *Tanichthys* leucophores had a pale-yellow hue of variable intensity among cells. Cyprinids thus have at least three leucophores: ML, from melanophores that sometimes contain residual melanin; XL, with white material and yellow/orange carotenoid-containing vesicles that can be mobilized by epinephrine; and *Tanichthys* leucophores, with white to pale yellow material alone.

To better understand these pattern features and their cytological bases, we examined cells by transmission electron microscopy. ML ultrastructure is well-documented (23, 25) so we focused on XL of zebrafish and leucophores of *Tanichthys*. Newly differentiated XL had abundant organelles that resembled pteridine-containing pterinosomes of xanthophores (Fig. 1*C*), though tending to be larger and less regular in shape than presumptive pterinosomes in yellow, xanthophore-like cells (∼X) also found in the anal fin at this early stage (see below). XL also exhibited small electron-dense organelles that resembled lipid containing vesicles, similar to carotenoid vesicles in xanthophores of the dorsal fin and body interstripes [e.g., (23, 26)]. Cryptic xanthophores (Xc) without overt pigmentation among fin melanophores had fewer, less developed pterinosomes and lacked lipid droplets.

Leucophores as well as xanthophores of *Tanichthys* had pterinosome-like organelles (Fig. 1*D*), which were larger and more electron-dense than those of XL and ∼X in zebrafish. Leucophore organelles also differed from those of red erythrophores, which harbored small presumptive pterinosomes and presumptive carotenoid vesicles with electron-dense peripheries. Thus, neither XL nor leucophores contained organelles with jagged outlines reminiscent of guanine-crystal containing organelles of zebrafish ML, nor did either cell exhibit organized reflecting platelets of crystalline guanine as observed in iridophores (28, 35), or vesicles with star-like crystals of uric acid observed in leucophores of *Oryzias* (30).

### XL Develop Progressively from Xanthophore-like Cells During Fin Outgrowth

To learn how XL populate the fin and their relationship to xanthophore like cells (∼X), we examined the ontogeny of XL appearance (Fig. 2*A*). XL were first evident among ∼X as cells with nascent accumulations of white material when the first (proximal) fin interstripe was forming. As development proceeded, additional XL were apparent until ∼X, lacking white material, were no longer evident. With continued fin outgrowth and appearance of a new distal interstripe populated by ∼X, a gradual increase in numbers of XL was evident, with correspondingly reduced numbers of ∼X. In the adult, interstripes were fully populated by XL, except for the most distal interstripe where some ∼X persisted; the appearance of small numbers of iridophores first near fin rays, and later between rays, contributed to the overall conspicuousness of the interstripes as well.

**Fig. 2.**
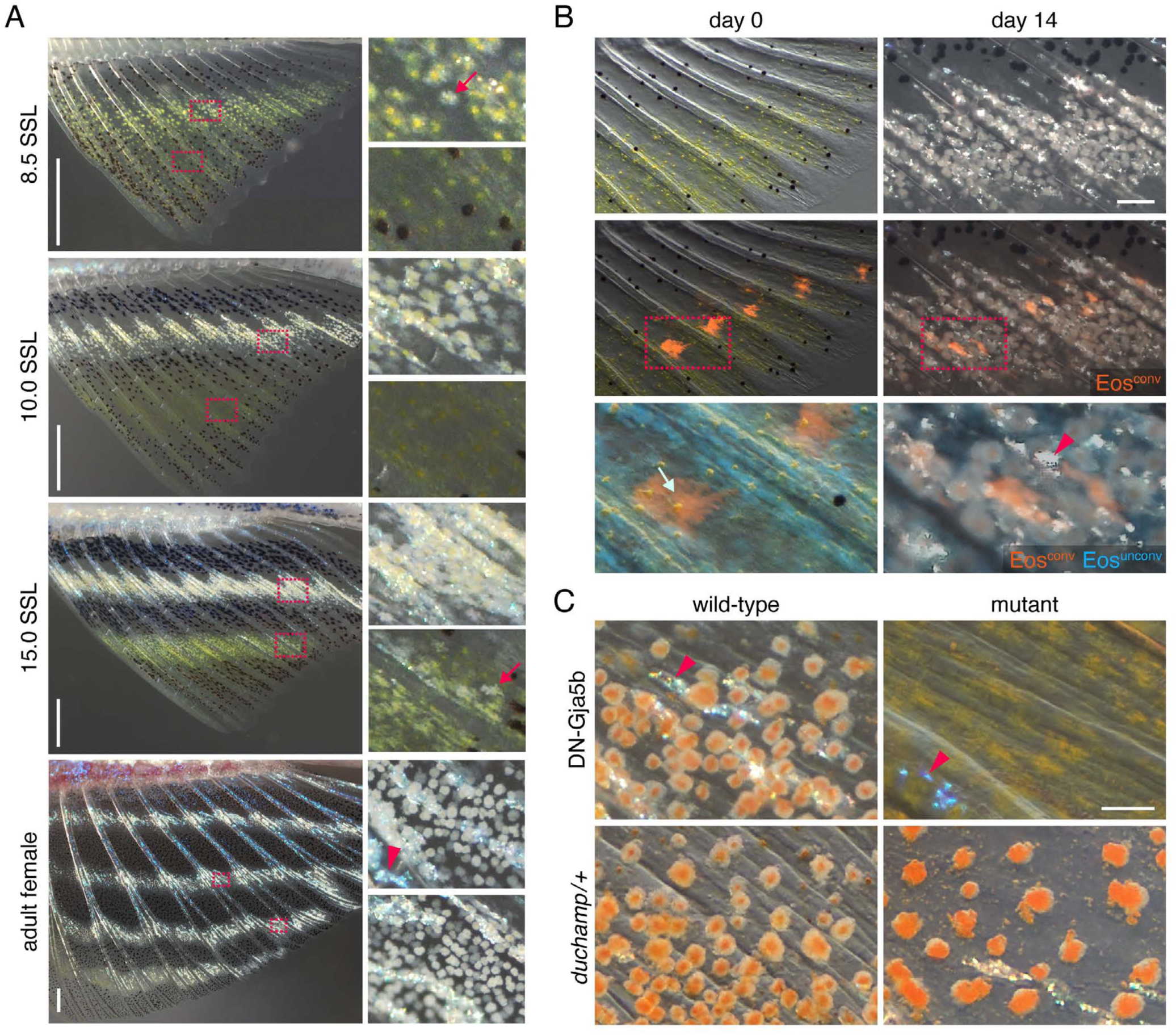
Conversion of ∼X cells to XL and genetic dependence on gap junctional communication and Aquaporin channel activity. (*A*) During fin outgrowth, XL appeared sequentially in regions where ∼X had already differentiated. Left panels show low magnification with higher magnifications of boxed regions at right. At 8.5 SSL cells with a white phenotype had just started to appear proximally whereas more recently differentiated ∼X were evident among melanophores distally. By 10 SSL, all proximal cells had an XL phenotype and ∼X were present in the nascent distal interstripe. At 15 SSL, XL appeared in the distal interstripe (arrow); within each interstripe XL appeared posteriorly and were gradually found more anteriorly. In the adult, XL were present in all fin interstripes and iridescent iridophores (e.g., arrowhead) had also differentiated at these sites. (*B*) ∼X acquired the XL phenotype by 14 d after photoconversion of *mitfa*:Eos at ∼8.0 SSL. Boxed regions shown at higher magnification in lower panels (arrow on d0, pigment granules contracted by epinephrine treatment). (*N*=6 fish, 27 xanthophores converted at day 0 leading to 37 marked XL at day 14, with cell number increase indicative of on-going proliferation; iridophore iridescence marked with arrowhead). (*C*) ∼X failed to transition to XL in fish homozygous for a dominant negative allele of Gja5b whereas XL had reduced white material and irregular morphologies in mutants heterozygous for *duchamp*, encoding the water, glycerol and peroxide channel Aquaporin 3a. Mutants are shown with wild-type siblings. (Scale bars: *A* 500 µm, *B* 100 µm, *C* 50 µm).

The sequence of XL appearance and ∼X disappearance suggested a direct transition. To test this model, and exclude the possibility that ∼X die and are replaced by XL from an unpigmented precursor, we fate-mapped ∼X by photoconverting Eos fluorescent protein driven by regulatory elements of *micropthalmia a* (*mitfa*) (36). When we marked ∼X of the nascent proximal interstripe in this manner, these cells or their progeny persisted and acquired a white XL phenotype (Fig. 2*B*). Thus, ∼X transition directly to XL, consistent with a requirement by XL for “xanthogenic” signaling and transcription factor activity via Colony Stimulating Factor-1 Receptor and Mitfa, respectively (23). This transition was reminiscent of M–ML transition in the dorsal fin, which is stimulated by overlapping signals that confer positional information (25). Yet the ∼X–XL transition occurred in a context that resembled the reiterative formation of interstripes on the body, which depends on self-organizing interactions among pigment cells in an expanding domain as the fish grows (37, 38).

### Zebrafish Pigment Pattern Mutants Reveal Genes Required for XL Differentiation

The spatial and temporal dynamics of ∼X–XL transition led us to ask if mechanisms that drive interactions among pigment cells on the body are also required by XL in the fin. Likely the best known contributor to body stripes is communication within and between cell types mediated by gap junctions, dependent on connexins encoded by *gja5b* (*leopard*) and *cx39.4* (*luchs*); if either gene is mutated a pattern develops of spots instead of stripes (39, 40). To see if such communication promotes development of XL we examined fish mutant for dominant negative allele *gja5b^stl710^* (amino acid substitution E42K), which blocks current flow across gap junctions that are either homomeric or heteromeric (Gja5b, Cx39.4) (41). Unlike wild-type siblings, XL of mutants lacked white material, retaining the appearance of ∼X progenitors (Fig. 2*C*; *SI Appendix*, Fig. S1).

We examined other pattern mutants and found two with spotted phenotypes, *duchamp* and *chagall*, in which XL lacked or had reduced white material, with evident expansions in yellow/orange carotenoid accumulations (Fig. 2*D*). We cloned these mutants and found both to be missense alleles of the transmembrane channel gene *aquaporin 3a* (*aqp3a*; amino acid substitutions A67D and R225C, respectively; *SI Appendix*, Fig. S2*A–E*,*G*). *aqp3a* was broadly expressed in fin, with transcript in both pigment cells and other cells of the tissue environment (*SI Appendix*, Fig. S2*F*). AQP3 is permeable to water, regulating water balance, but also glycerol and H_2_O_2_, and in this latter role it contributes to cellular redox state, with implications for progression of melanoma and other tumors in which the channel is upregulated (42, 43). We found that mutant proteins of both zebrafish alleles had grossly reduced permeabilities to all three solutes and dominant negative activities against wild-type channel, leading to defects in ∼X-XL of the fin though not xanthophores on the body (*SI Appendix*, Fig. S3).

### Enhanced Expression of Pteridine Genes in XL and Requirement for Sepiapterin Reductase

To further understand ∼X–XL transition, and pathway(s) required for the white phenotype, we compared gene expression by bulk mRNA sequencing of cells purified by flow sorting for *aox5*:memEGFP (23), from proximal (XL) and distal (∼X) fin tissues dissected separately at 15 SSL (Fig. 2*A*). Compared to ∼X, XL had more abundant transcripts for 1446 genes (log_2_ fold-change>1, *q*<0.05), of which at least 65 have been studied for roles in other pigment cell types (44)(Fig. 3*A*; *SI Appendix*, Fig. S4). Consistent with gap junction and Aqp3a requirements, XL had higher transcript abundances for *gja5b* (log_2_FC=1.06, *q*=0.002), *cx39.4* (log_2_FC=1.98, *q*=6.8E-24) and *aqp3a* (log_2_FC=0.85, *q*=1.2E-46) compared to ∼X (Dataset S1).

**Fig. 3.**
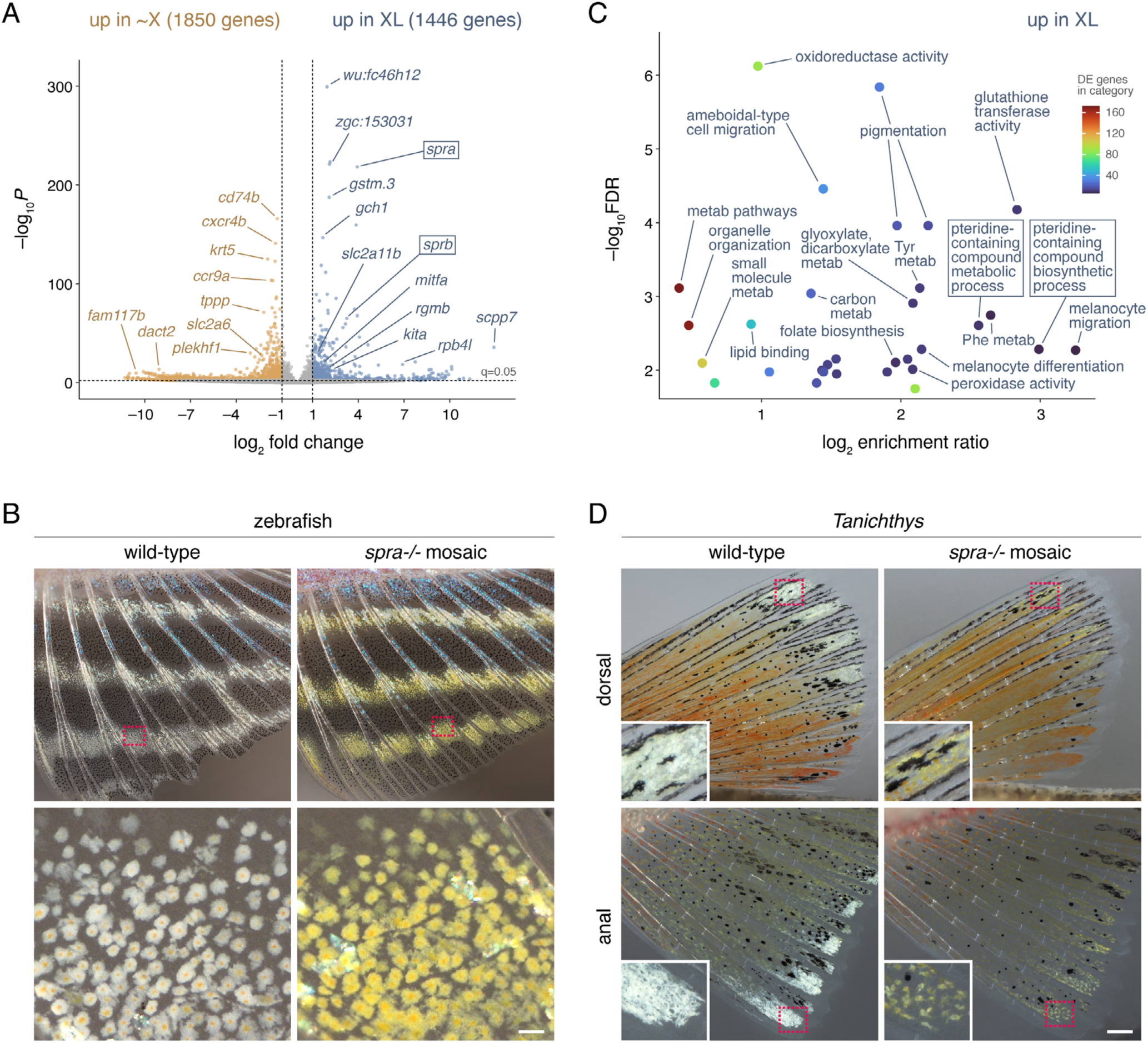
Transcriptomic comparison of ∼X and XL in zebrafish and requirement for sepiapterin reductase in acquisition of white phenotypes in zebrafish and *Tanichthys*. (*A*) Sepiapterin reductase genes *spra* and *sprb* were differentially expressed between ∼X and XL at 15 mm SL, as were numerous genes with previously described pigmentary roles. *mitfa and kita* were expressed at higher levels in XL than xanthophores: *mitfa* is required whereas *kita* is dispensable for XL development (23). *slc2a11b* is required for leucophore development in *Oryzias* (61). (*B*) Zebrafish mosaic for *spra-/-* mutations developed XL that retained the earlier yellow appearance of ∼X progenitors. (C) Leucophores of *Tanichthys* mosaic for *spra-/-* mutations had gross reductions in white material yet failed to develop yellow pigmentation to the same excess as XL. (*D*) Pathway enrichment for genes with higher transcript abundances in XL than ∼X showing combined analyses for gene ontology terms of biological and molecular functions as well as KEGG pathways. Enrichment for pteridine pathways as well as oxidoreductase and glutathione transferase activity were particularly notable. Terms that reference melanocytes often include genes that function in multiple chromatophore types of zebrafish. (FDR, false discovery rate). (Scale bars: *C* 500 µm, *D* 200 µm).

Notable, too, was higher expression in XL of *sepiapterin reductase a* (*spra*) and *sprb* (Fig. 3*A*), encoding enzymes of the pteridine pathway. Sepiapterin is a yellow pigment responsible for the color of embryonic xanthophores in zebrafish but several other pteridines are pale, white or colorless (45–48), raising the possibility that sepiapterin reductase contributes to an accumulation of white or colorless pteridines in XL. To test this idea, we targeted coding sequences of *spra* and *sprb* by CRISPR/Cas9 mutagenesis. We observed no pigmentary phenotype in genetically mosaic fish injected with reagents targeting *sprb*. By contrast, somatic mutagenesis of *spra* resulted in the loss of white material and a prominent yellow color in XL (Fig. 3*B*; *SI Appendix*, Fig. S5). Fish homozygous for mutations in *spra* were larval-lethal and so not informative for this pigmentary trait.

Given presumptive pterinosomes in leucophores of *Tanichthys*, we hypothesized that sepiapterin reductase-dependent pteridines contribute to white fin ornamentation of this species as well. We sequenced the genome, identified an orthologue of *spra*, and targeted the locus by CRISPR/Cas9 mutagenesis. Somatic mutations led to markedly reduced white material in leucophores of dorsal and anal fins, but without a correspondingly dramatic increase in yellow color (Fig. 3*D*). Thus, leucophores in both locations of *Tanichthys* resembled XL of the zebrafish anal fin in requiring *spra*. But the failure of *spra* mutation to drive yellow pigment overabundance in *Tanichthys* suggests that pale or colorless pigments derived from sepiapterin reductase activity may arise from different precursors in this species, or that homeostatic mechanisms different from those of zebrafish check yellow pigment production in the absence of sepiapterin reductase activity.

The *spra*-dependence of pigmentary phenotype led us to assess more comprehensively if additional genes of this pathway were modulated in zebrafish XL relative to ∼X. Gene enrichment analyses confirmed upregulation in XL of pathways for pteridine-containing compound metabolism and biosynthesis (Fig. 3*C*; *SI Appendix*, Fig. S4; Dataset S2). Genes of other pathways with strong signatures encoded products with oxidoreductase, glutathione transferase and lipid binding activities. In xanthophores of the body, lipid accumulation is associated with terminal differentiation and carotenoid deposition (26). Consistent with enhanced lipid binding activity in XL, we observed an accumulation of Oil-red-O+ lipid droplets in XL compared to ∼X (*SI Appendix*, Fig. S6).

### Pathway Analyses of XL Indicate Conditions for Generating Pale or Colorless Pteridines

Pteridines comprise fused pyrimidine and pyrazine rings that occur in pterins (pteridines with lactam and amino groups) like sepiapterin, but also many other compounds including the enzymatic cofactor tetrahydrobiopterin (H_4_-Biopterin) and its precursors and derivatives, and the co-enzyme folic acid (*SI Appendix,* Fig. S7*A*) (46–48). Pteridine-related gene sets revealed by RNA-seq included several loci encoding enzymes in this pathway (Fig. 4*A*; Dataset S3): in addition to sepiapterin reductases, encoded by *spra* and *sprb* (steps 3, 5, 15 in Fig. 4*A*), these included GTP cyclohydrolases of *gch1* and *gch2* (step 1), carbonyl reductases of *cbr1* and *cbr1l* (step 4), quinoid dihydropteridine reductase of *qdpra* (step 9), xanthine dehydrogenase of *xdh* (steps 17, 21, 25), and dihydrofolate reductase, an activity predicted for *zgc:153031* (step 16). Additional factors were phenylalanine hydroxylase of *pah* that requires H_4_-biopterin cofactor (step 7), aldehyde oxidase of *aox5* (strong candidate for step 18), and MYC binding protein 2 of *mycbp2* (*esrom*), an E3 ubiquitin ligase required for yellow sepiapterin production in embryonic xanthophores (45), likely via high-level pathway regulation. These observations and results of *spra-/-* mutagenesis suggested an abundance of pteridines in XL, an inference supported by ammonia-induced autofluorescence typical of pteridines (*SI Appendix,* Fig. S7*B*).

**Fig. 4.**
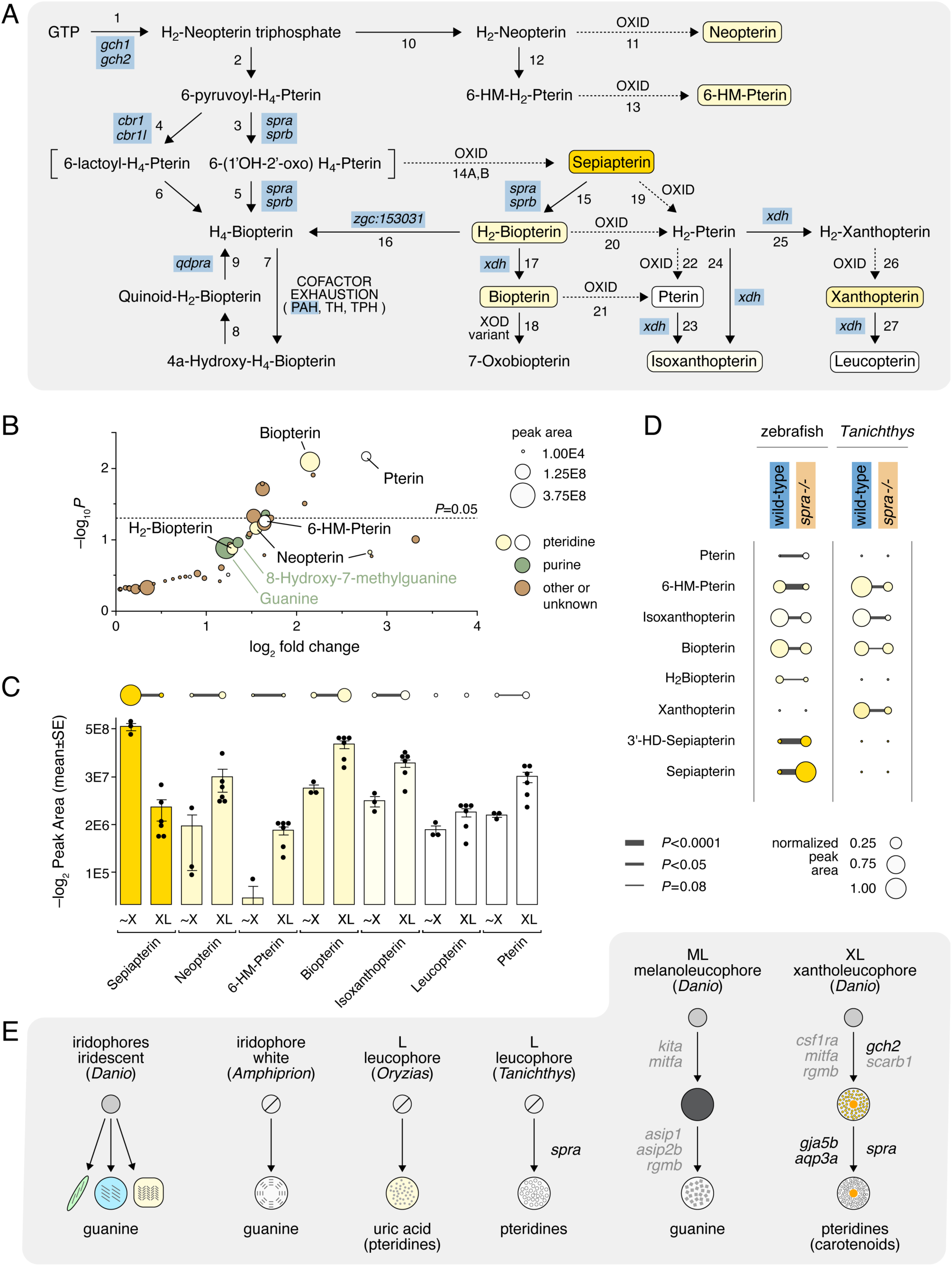
Chemical analyses point to pale and colorless pteridine accumulation by zebrafish XL and *Tanichthys* leucophores. (*A*) Pteridine pathway showing compounds identified here (outlined) and intermediates, as well as genes with higher transcript abundance in XL than ∼X and enzymatic activities corresponding to specific pathway steps (blue boxes). [Structure after (46–48) as well as KEGG pathway maps and notations provided in Dataset S3.] (*B*) Compounds with greater abundance in XL+ tissue than XL– tissue after controlling for effects of anatomical position and genotype in metabolomic analyses (*P*-values shown for one-tailed tests under hypothesis of abundance in XL+ >> XL–). XL+ tissue had more abundant biopterin and pterin. (*C*) Diminished yellow pteridines and increased pale or colorless pteridines in XL+ tissue compared to tissue containing ∼X assayed by targeted mass spectroscopy. Bar plot shows log_2_-transformed values for visualizing within-pteridine differences and dots above show non-transformed amounts for visualizing relative abundances across pteridines, normalized to the most abundant type (sepiapterin in ∼X); horizontal line widths indicate significance levels of pairwise comparisons. (*D*) Metabolomic analyses of zebrafish and *Tanichthys* mosaic for *spra* mutations showing reductions in pale or colorless pteridines compared to wild-types, with zebrafish *spra-/-* mosaics also exhibiting increased quantities of yellow pteridines; significance levels illustrated as in *C*. (*E*) Cell lineages, chemical and structural bases, and genetic requirements for white cell phenotypes within and between species. Small grey circles denote progenitors identified by fate mapping, small circles with slashes indicate lineages not fully defined. See text for additional details.

Enhanced pteridine pathway gene expression in XL suggested that mutations upstream of *spra* should also impact XL pigmentary phenotype. We therefore examined XL in fish homozygous mutant for a splicing-defective allele of *gch2* (49) (Fig. 4*A*, step 1). XL in such individuals had reduced white material but also lacked the striking yellow phenotype of XL in *spra-/-* mosaics, consistent with diminished pteridine contents overall (*SI Appendix,* Fig. S8*A*).

Several reactions in the pteridine pathway are oxidations, some mediated enzymatically (e.g., Fig. 4*A*, steps 17, 18), but others thought to be spontaneous (steps 11, 13, 14A, 14B, 19–21, 26). Substantial upregulation of genes encoding oxidoreductases and glutathione transferases (Fig. 3*C*) suggested oxidative conditions. as did greater abundances of transcripts specifically encoding enzymes for oxidative stress response and peroxide detoxification, including peroxiredoxins 1, 2 and 6 (log_2_FC=0.9–1.1, *q*<7.4E-7), catalase (log_2_FC=0.7, *q*<0.002), NAD(P)H dehydrogenase, quinone 1 (Nqo1) (log_2_FC=1.2, *q*<7.7E-7) and thioredoxin (log_2_FC=0.75, *q*<1.7E-5) (Dataset S2) (50–52). These observations, and high levels of sepiapterin reductase and xanthine dehydrogenase gene expression, suggested that XL drive pteridine production from yellow sepiapterin towards pale or colorless pteridines.

### Chemical Analyses of White-Cell Containing Tissues Indicate Pale and Colorless Pteridine Accumulation in XL and Leucophores

To determine if XL are specifically enriched for pale or colorless pteridines relative to other cell types, we used a metabolomic approach. We hypothesized that pteridines contributing white material to XL should be markedly more abundant (log_2_FC>1) in (proximal) tissue harboring XL at 14 SSL compared to tissues without these cells. Yet we also anticipated that analyses could be confounded by iridophores that co-occur with XL (e.g., Fig. 2*A*) or by proximal–distal differences unrelated to pigment cells. To remove confounding effects of iridophores we used fish that lacked these cells, owing to a mutation in *leukocyte tyrosine kinase* (*ltk*) (53). To control for proximal–distal differences, we engineered these *ltk-/-* fish to have XL either in the normal or displaced distally, owing to a recessive mutation in *rgmb*, which encodes a non-canonical BMP receptor (25, 54). We could therefore compare tissues containing XL (XL+) to tissues without XL (XL–), taking a multifactorial approach to control for variation between anatomical regions and *rgmb* genotypes. This analysis revealed 12 compounds significantly enriched in XL+ tissue, including biopterin and pterin, along with additional pteridines that tended towards excess; all these pteridines are pale-yellow or colorless (Fig. 4*B*; *SI Appendix* Figs. S7*A* and S8*C*; Datasets S4–S6). Also evident at levels of enrichment that failed to reach statistical significance were several purines; these included guanine, but also 8-Hydroxy-7-methyguanine, consistent with oxidative conditions (Fig. 4*B*; *SI Appendix*, Fig. S8*C*) (55).

Given these initial findings for XL, and the yellow color of ∼X progenitors, we hypothesized that sepiapterin is produced in ∼X and converted to pale or colorless forms during XL differentiation. Accordingly, we predicted reciprocal abundances of these forms in comparisons of ∼X+ and XL+ tissues. Targeted mass spectroscopy confirmed higher quantities of yellow pteridines in ∼X+ tissue and higher quantities of pale yellow to colorless pteridines in XL+ tissue (Fig. 4*C*; Dataset S7). As an independent test of this prediction using wild-type fish (*ltk*+/+, *rgmb*+/+) at the same stage, we undertook a metabolomic comparison, which confirmed higher abundances of yellow vs. pale or colorless pteridines in ∼X+ and XL+ tissues, respectively (*SI Appendix*, Fig. S8*D*; Datasets S8 and S9).

If pale or colorless pteridines are responsible for the phenotypes of both zebrafish XL and *Tanichthys* leucophores, then *spra-/-* mutations that diminish white material (Fig. 3*B*,*D*) should reduce the quantities of these pteridines. Confirming this prediction, metabolomic analyses revealed significant reductions in pale and colorless pteridines in *spra*-/- mosaic fins compared to wild-type fins (Fig. 4*D*; Datasets S10–S13). In zebrafish, these changes were accompanied by increased quantities of yellow sepiapterin and 3’ hydroxy-D-sepiapterin. By contrast, increases in colored pteridines were not evident in *Tanichthys* mosaic for *spra* mutations. Unexpectedly these analyses also indicated a significant difference in guanine abundance between wild-type and mosaic *spra-/-* mutant tissue of *Tanichthys* (*SI Appendix*, Fig. 8*F*). This result prompted us to assess guanine as well as uric acid quantities in leucophore-containing vs. other fin tissue of *Tanichthys* by targeted mass spectroscopy. These comparisons indicated moderately greater amounts of guanine, though not urate, in tissue with leucophores, despite the absence of morphological evidence for guanine crystals by TEM (*SI Appendix*, Fig. 8*E*, Fig. 1*D*; Dataset S14).

## Discussion

Melanocytes and chromatophores are a well-researched system for elucidating mechanisms of fate specification in neural crest lineages. The most-studied chromatophores—melanophores, xanthophores and iridophores—have provided insights into gene regulatory networks for specification and differentiation (53, 56), and more recently, how requirements for genes have evolved across species (36, 57, 58). In this study we have elucidated features of white chromatophores, uncovering a remarkable diversity in cellular and biochemical phenotypes, with apparently independent evolutionary origins—across neural crest cell sublineages and across teleost phylogenetic lineages.

These and prior efforts indicate at least six classes of highly reflective chromatophores (Fig. 4*E*). Iridophores consist of several subtypes, even in zebrafish, that vary in shape, density, anatomical location, guanine reflecting platelet sizes and arrangements, and hues reflected (28, 59). Beyond zebrafish are other iridophore morphologies, including those of anemonefish, *Amphiprion*, in which a different organization of guanine reflecting platelets leads to a white rather than iridescent phenotype (60). A white phenotype is achieved in a different way in leucophores of *Ozyzias*, relying on crystals of uric acid (29, 30). The lineage of *Oryzias* leucophores is not certain yet embryonic leucophores are thought to have affinity to xanthophores (56, 61) and adult leucophores to resemble xanthophores in some respects and iridophores in others (30).

Within cyprinid fishes, ML are white because of irregular guanine crystals. By contrast, we found that leucophores of *Tanichthys*, though similar in some respects to ML, were enriched for pale or colorless pteridines, with pteridine complements and the white phenotype itself depending on sepiapterin reductase. Similarly for XL of zebrafish, an essential role for pale or colorless pteridines in generating the “off-white” component of these cells was suggested by the greater abundance of these compounds in chemical analyses of XL-containing tissue, and the conversion of a white to yellow phenotype, with associated changes in pteridine composition, in mosaics for *spra* mutation. Though pteridines such as those identified here—including pterin and leucopterin in XL, and isoxanthopterin of *Tanichthys* leucophores—are not typically associated with pigmentary roles in vertebrates, they have been studied for contributions to invertebrate integuments (48, 62, 63). Likewise in insects, sepiapterin reductase mutants of silkworm are bright yellow due to sepiapterin accumulation, though a normally white integument in this species results from xanthine dehydrogenase dependent uric acid deposition (64–66). Although our analyses implicate pale or colorless pteridines in XL and leucophore phenotypes, we cannot exclude contributions from other chemicals or specific structural features of these cells, whether specializations in pteridine organization, light scattering lipid droplets, or purines in non-crystalline yet somehow reflective forms (21, 23, 67, 68). Resolving photonic properties of these cells, and how a white phenotype is achieved with pteridines that are pale or colorless in solution and perhaps other compounds will be interesting to uncover, as has recently been achieved through elegant chemical and structural analyses of *Oryzias* leucophores and other reflective cells (30, 35, 59).

Our several approaches also suggest a model for development of XL with parallels and differences compared to ML. Both arise from already-pigmented cells: melanophore progenitors of ML resemble other melanophores whereas ∼X that give rise to XL lack the concentrations of carotenoids found in adult xanthophores of the body, and appear to derive their yellow color principally from sepiapterin, much like embryonic xanthophores (26, 45). M–ML transition requires high levels of intersecting BMP and Agouti signals, with cellular remodeling that involves demelanization by an autophagous process and production of new guanine-crystal containing organelles (25). By contrast, ∼X–XL transition occurs at low levels of BMP signaling and does so dynamically as fin interstripes form and the fin grows, continuing even in *rgmb* mutants in which XL are displaced distally (54). This transition also requires gap junctional communication, similar to chromatophore patterning of stripes on the body, but unlike differentiation of ML. As X–XL transition proceeds, cells upregulate several pteridine synthesis genes, especially *spra* and *sprb* the products of which presumably generate H_4_-biopterin while also converting yellow sepiapterin to pale H_2_-biopterin. Concomitantly enhanced expression of *xdh* allows further conversions towards pale or colorless pteridines, as does an intracellular redox state supportive of spontaneous oxidations in the same direction, inferred from transcripts for oxidative stress and detoxification genes, oxidized purine, and the abundance of pterin itself, thought to arise by spontaneous oxidation. In this context, the white-deficient phenotype of mutants for Aqp3b is intriguing, given roles for this channel in peroxide transport and modulation of redox conditions. Thus, although ML and XL are similar in having a white phenotype, the cell types differ markedly in sublineage origins, remodeling, and chemical bases of pigmentary material. It will be interesting to learn what other factors might trigger X–XL transition and how physiological states enabled by gap junctional communication and Aqp3a activity allow this transition to proceed.

White cells are found across a variety of teleosts yet the cell types described here are not so common phylogenetically as to have arisen from a common ancestral white cell, nor would their molecular, cell lineage, structural, or chemical features support a single evolutionary origin. Rather, these cells appear to have evolved independently across neural crest sublineages and phylogenetic lineages, with properties either convergent or parallel depending on comparison and level of analysis. They present an outstanding model in which to identify genomic, gene regulatory and developmental mechanisms underlying the evolution of novel cell types. The shared derivation of these cells from neural crest suggests further comparisons should provide useful insights into how such types have evolved, much as comparisons of closely related species or divergent populations have provided evolutionary genomic insights into diversification at the organismal level.

## Materials and Methods

### Animal husbandry

Zebrafish and *Tanichthys* were reared at ∼28 °C (14L:10D) and fed rotifers, Artemia and flake food. Fish were anesthetized in MS222 and in most instances treated with epinephrine to contract pigment granules towards prior to imaging.

### Mutagenesis and cloning

Mutations were induced by ENU or CRISPR/Cas9 using IDT AltR reagents and analyzed after isolation by standard genetic methods or as F0 mosaics. ENU-induced alleles were identified by standard methods of meiotic mapping or association mapping with pooled DNA, followed by verification of candidate lesions by functional assay as well as as induction of second-site mutations for phenotype reversion.

### Ultrastructural and Chemical Analysis

Cytological characteristics were assessed by transmission electron microscopy and chemical analyses performed by mass spectroscopy on dissected tissues.

### RNA-sequencing

Cells were collected by fluorescence activated cell sorting and replicate libraries prepared for sequencing, mapped by Kallisto and analyzed with DEseq2. Gene set enrichment analyses were performed using on-line tools at webgestalt.org.

### Data Sharing

Transcriptomic and chemical analyses are provided in Datasets S1–14. Transcriptomic data are accessible through NCBI GEO (Accession # GSE313210).

## ACKNOWLEDGEMENTS

We thank members of the Parichy lab for assistance with fish rearing, and Sarah Wilmsen and the UVA Molecular Electron Microscopy Core for assistance with electron microscopy. Supported by NIH R35 GM122471 (to D.M.P.), with support for Aqp3 channel analyses from Spanish Ministry of Science, Innovation and Universities (MICIU/AEI/10.13039/501100011033) and the European Regional Development Fund (European Union) Grant no. PID2022-138066OB-I00 to J.C. Thanks to L. B. Patterson for imaging *chagall* and *duchamp* and C.W. Higdon for assistance with meiotic mapping of *duchamp*.

## Supplementary Information

**Fig. S1.**
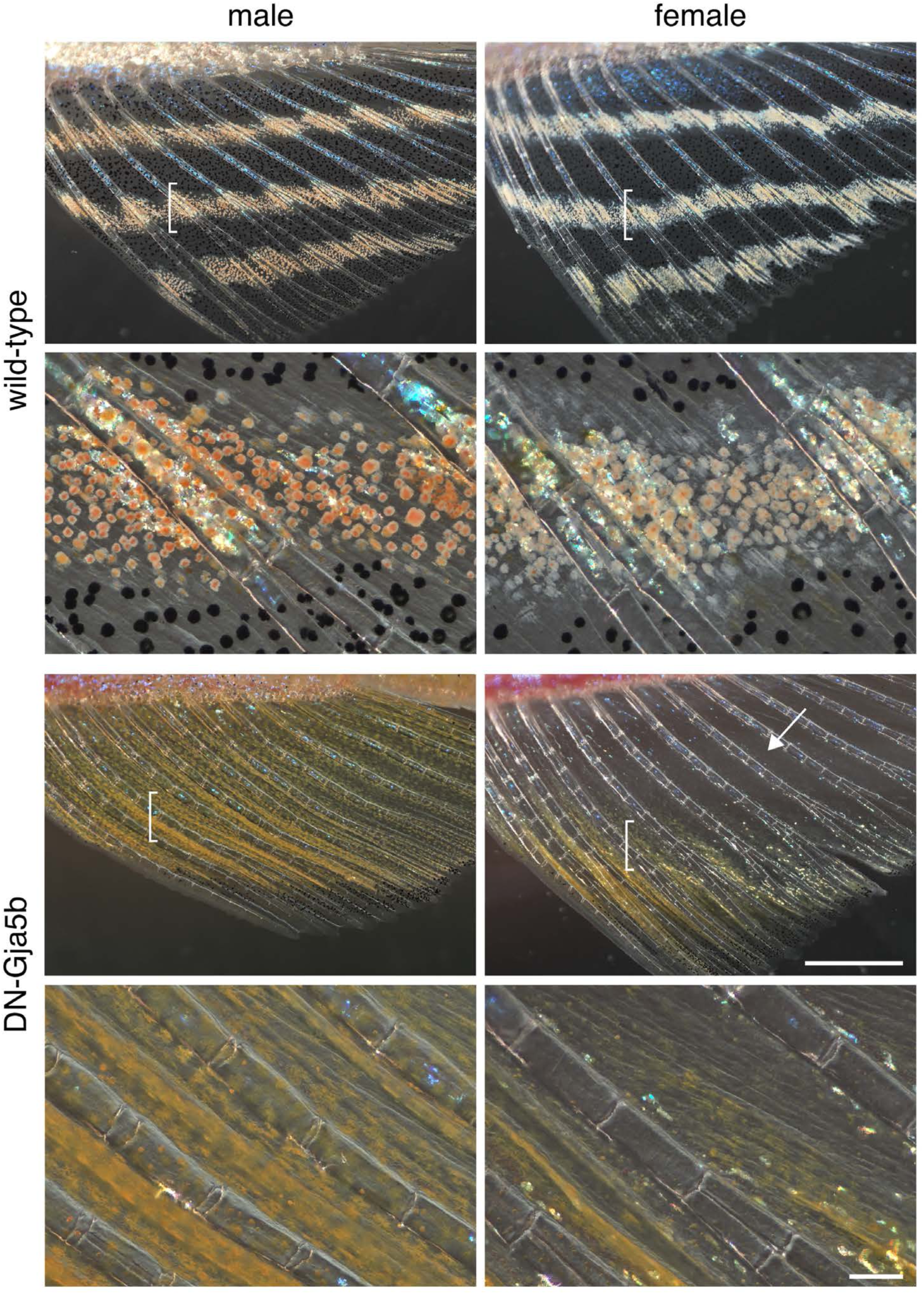
Phenotypes of dominant negative Gja5b allele *gja5b^stl710^* and differences in XL complements between males and females. In wild-type fish, XL of males are typically more numerous and brightly colored, with more prominent accumulations of orange carotenoids at cell centers after epinephrine treatment (1). In mutants for *gja5b^stl710^*, X-XL of males and females lacked white deposits and exhibited a relatively uniform yellow color with pigment granules that were not responsive to epinephrine treatment. Besides alterations in pigment phenotype, XL were markedly fewer in females than males, with a proximal region of fin (arrow) devoid of pigmented cells. Bracketed regions in upper panels are shown at higher magnification in lower panels. Homozygous mutants are shown; heterozygotes had intermediate phenotypes. (Scale bars: low magnification, 1 mm, high magnification 100 µm).

**Fig. S2.**
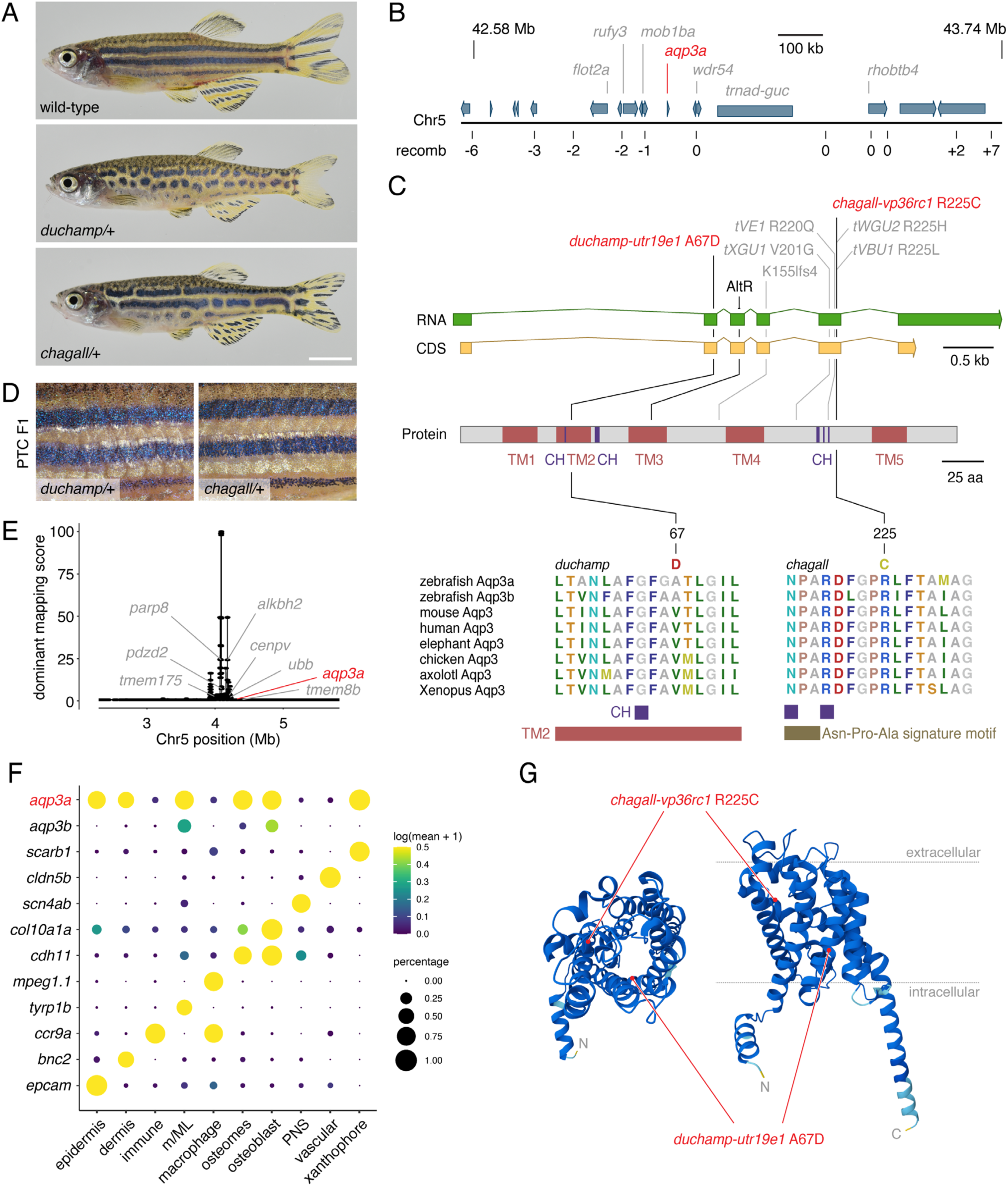
*duchamp* and *chagall* are alleles of *aqp3a*. (*A*) Heterozygous mutants are spotted and have fins somewhat shorter than wild-type; homozygotes have severe growth and fin defects and were not informative for XL phenotypes. (*B* and *E*) Meiotic mapping of ∼2000 individuals in B and association mapping of pooled phenotypes by whole-genome resequencing in *E*. Both approaches placed mutants in the vicinity of *aqp3a*, previously identified for pigment pattern defects of *mau* mutants (2). Only a subset of gene names is shown; *trnad-guc*, array of tRNA-Asp genes. (*C*) Gene body of *aqp3a* and protein structure showing locations of *duchamp* and *chagall* missense substitutions, alleles of *mau* (2) in grey, and the site targeted for CRISPR-Cas9 mutagenesis indicated (“AltR”). CH, amphipathic channel domain; TM1–5, transmembrane domains; Asn-Pro-Ala signature motif typical of aquaporin channels. (*D*) Confirming allelism of mutant phenotypes with *aqp3a* and suggesting dominant negative or neomorphic activities, targeting of mutant alleles to generate premature termination codons in the same background led to phenotypes indistinguishable from wild-type. (*F*) Dotplot of *aqp3a* and cell-type marker gene expression in reanalysis of single cell RNA-seq data of dorsal fins collected in (3). *aqp3a* was broadly expressed in several cell types, including xanthophores, and has been shown to have non-autonomous effects to pigment cells on the body (2). The paralogous gene, *aqp3b*, had a more limited expression domain. (*G*) Predicted structure of Aqp3a monomer (AlphaFold AF-D3TI80-F1-v4) with mutations indicated. Left structure viewed from extracellular face, right structure as oriented in membrane.

**Fig. S3.**
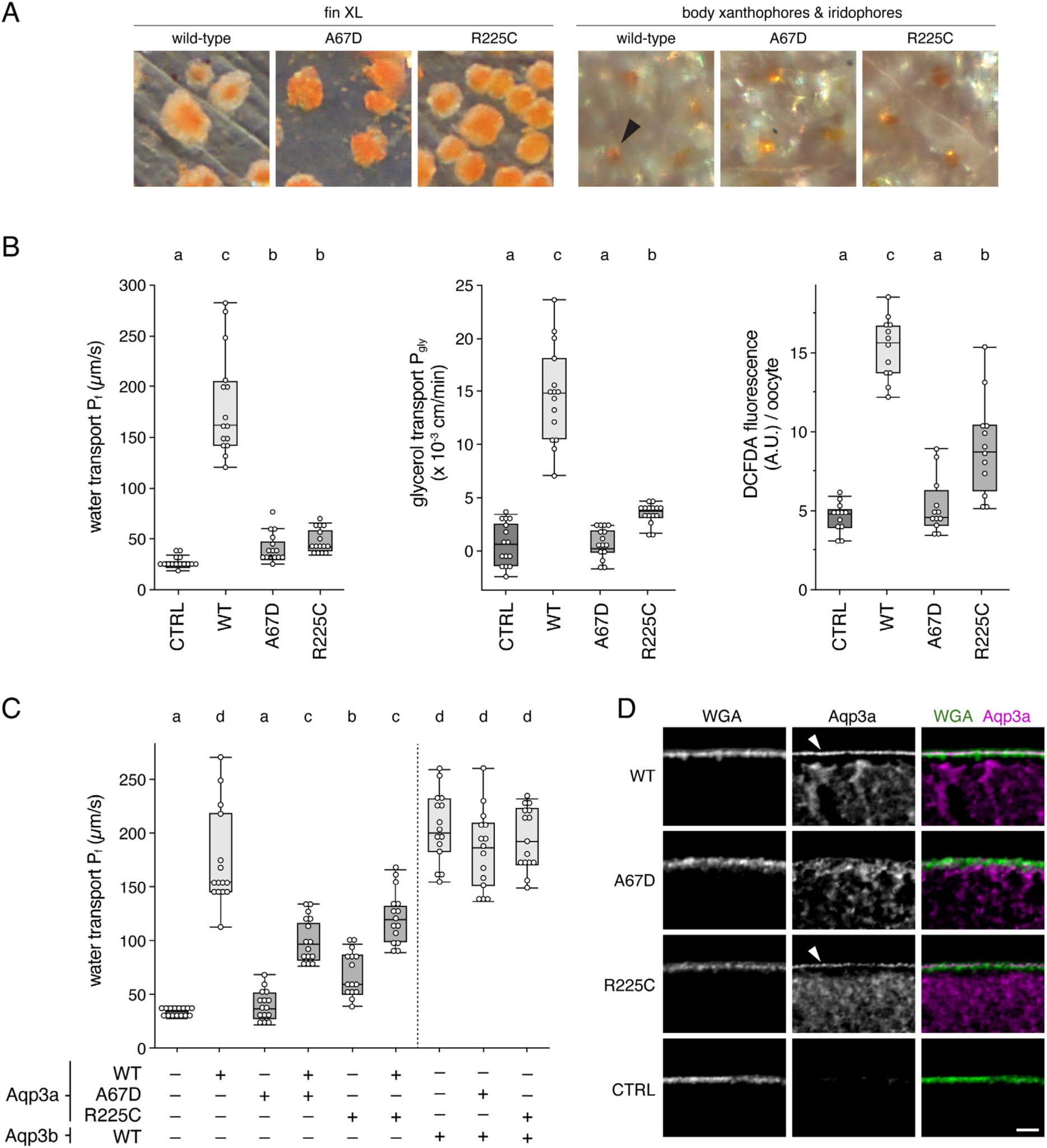
Aqp3a phenotypes and activities in *Xenopus* oocyte assays. (*A*) XL of fin across wild-type and heterozygous Aqp3a alleles (*duchamp*, *chagall*). Mutants carrying Aqp3a A67D (*duchamp*) were especially compromised for white material of XL and carriers of R225C (*chagall*) variably so. In both mutants, accumulations of yellow/orange carotenoids tended to be larger than those of wild-type. On the body, xanthophores had normal pigmentation (upper right panels; arrowhead, carotenoid vesicles of xanthophore aggregated by epinephrine). Nevertheless, embryonic, initially pteridine-pigmented xanthophores on the body of another *aqp3a* mutant allele are reported to lose contacts with one another and die during the larva-to-adult transition (2). Iridescent iridophores occur in a deeper plane of the skin, beneath xanthophores (4), and also appeared normal. (*B*) Transport of water, glycerol and peroxide was markedly reduced for A67D and R225C Aqp3a. Letters above groups indicate means not significantly different in Steel-Dwass *post hoc* comparisons (*P*>0.05; experiment-wide Kruskal-Wallis tests, all *P*<0.0001). (*C*) Tests of dominant negative activities for mutant Aqp3a against wild-type Aqp3a or the closely related product paralogous locus, Aqp3b. Left, Co-expression of A67D or R225C with wild-type Aqp3a significantly attenuated permeability to water, demonstrating dominant negative interactions. Right, Neither Aqp3a mutant impacted permeability through Aqp3b, consistent with lack of dominant negative activity against the product of its locus. (*D*) Immunofluorescence for Aqp3a revealed localization in Wheat Germ Agglutinin (WGA) labeled membrane for WT and R225C but not A67D. (Scale bar, 50 µm).

**Fig. S4.**
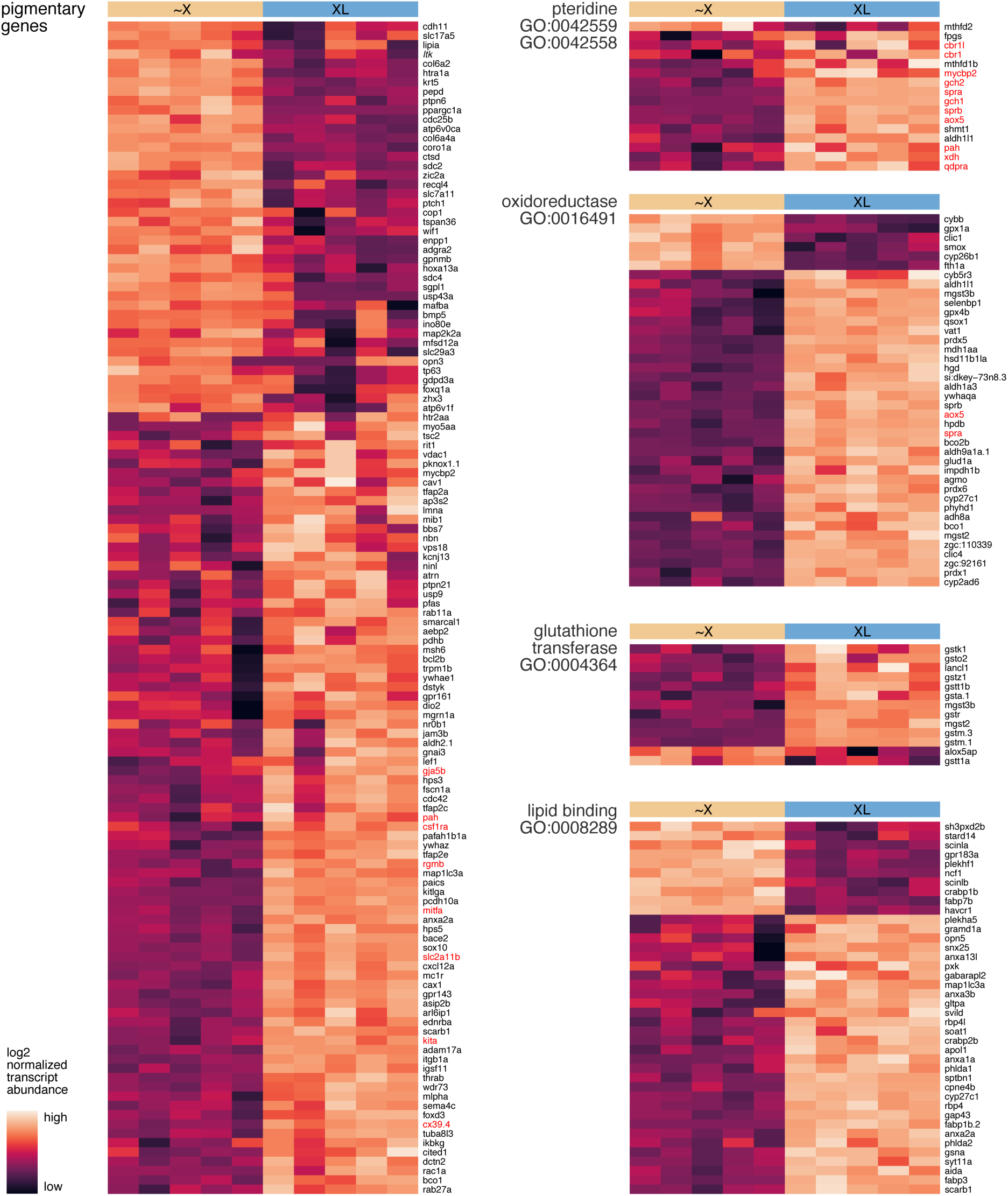
Heat maps of pigmentary gene (5) expression in ∼X and XL as well as genes associated with pteridine biosynthesis and metabolism, oxidoreductase activity (top 40 of 89 by *P*-value), glutathione activity and lipid binding (top 40 of 53). Some genes appear in more than one category and several genes mentioned in main text are shown in red. Each column represents expression in a replicate ∼X or XL library.

**Fig. S5.**
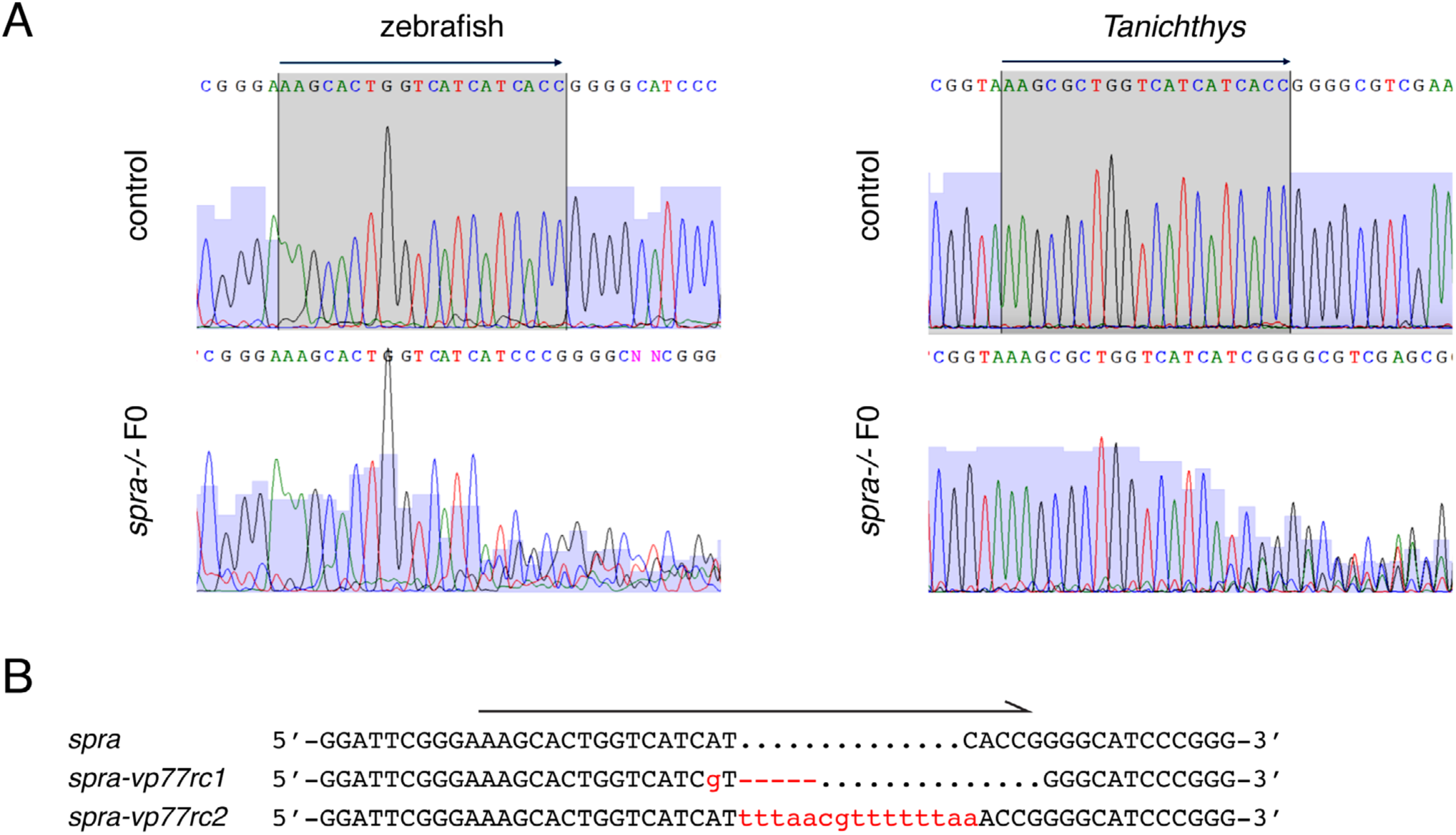
Mutations in *spra*. (*A*) Example electropherograms for wild-type control and mosaic (F0) *spra-/-* mutant embryos of zebrafish and *Tanichthys*. Shaded regions and arrows above indicate target sites. (*B*) Mutations recovered in F1 *spra* mutant zebrafish (-5 bp, +14 bp) that were larval-lethal (e.g., 38 sibling adults genotyped by Sanger sequencing yielded 13 homozygous wild-type, 25 heterozygotes and 0 homozygous mutants).

**Fig. S6.**
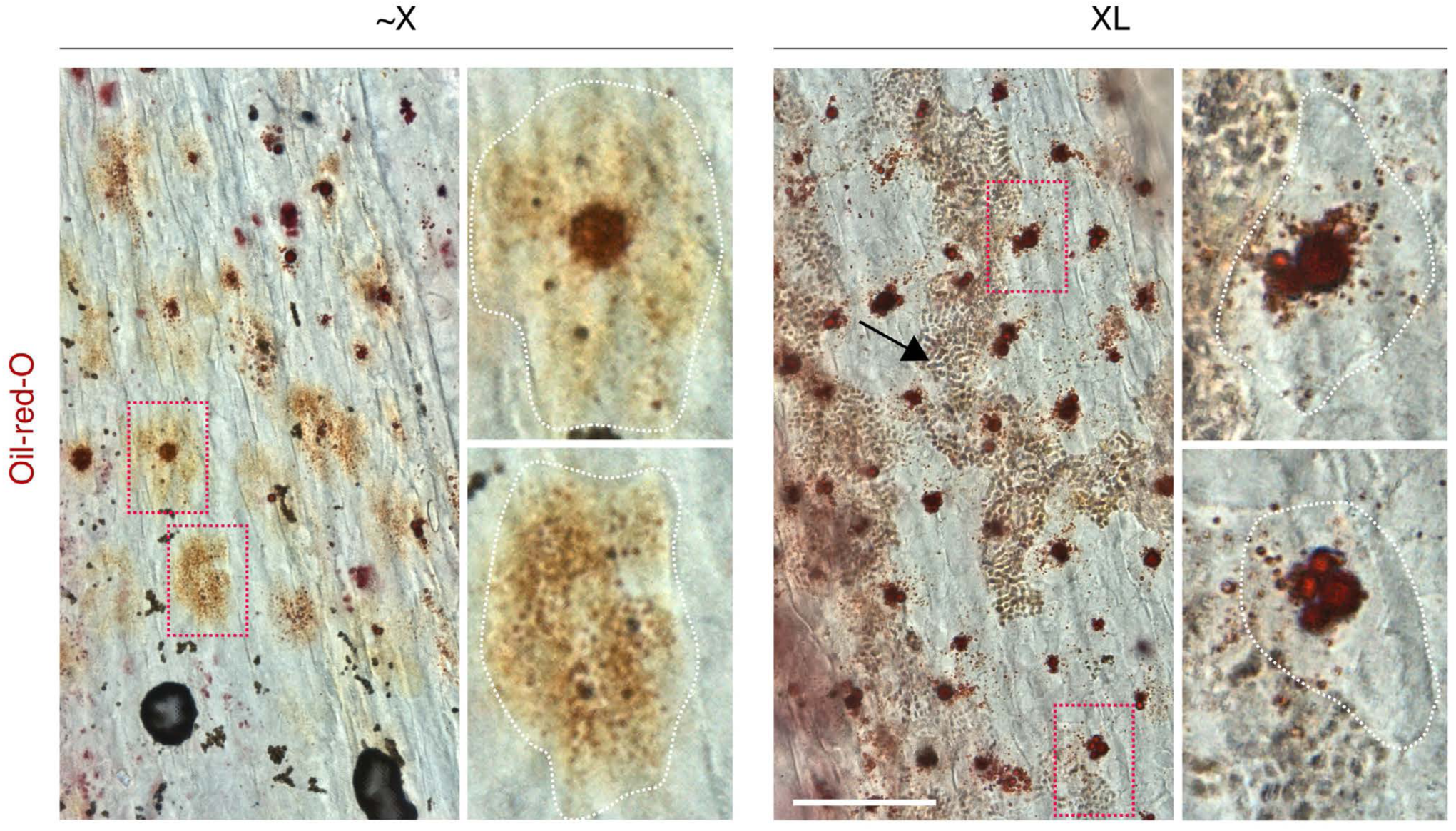
Lipid droplet accumulation in XL revealed by Oil-red-O staining. ∼X had variable levels of Oil-red-O staining consistent with early stages of carotenoid vesicle production. XL had prominently stained droplets concentrated towards cell centers on epinephrine treatment, where carotenoid vesicles are found, though some droplets remained relatively dispersed. Regions of dotted red boxes shown at higher magnification. Approximate cellular boundaries indicated with dotted white lines. Arrow in XL panel indicates reflecting platelets of co-occurring iridophores. (Scale bar, 100 µm).

**Fig. S7.**
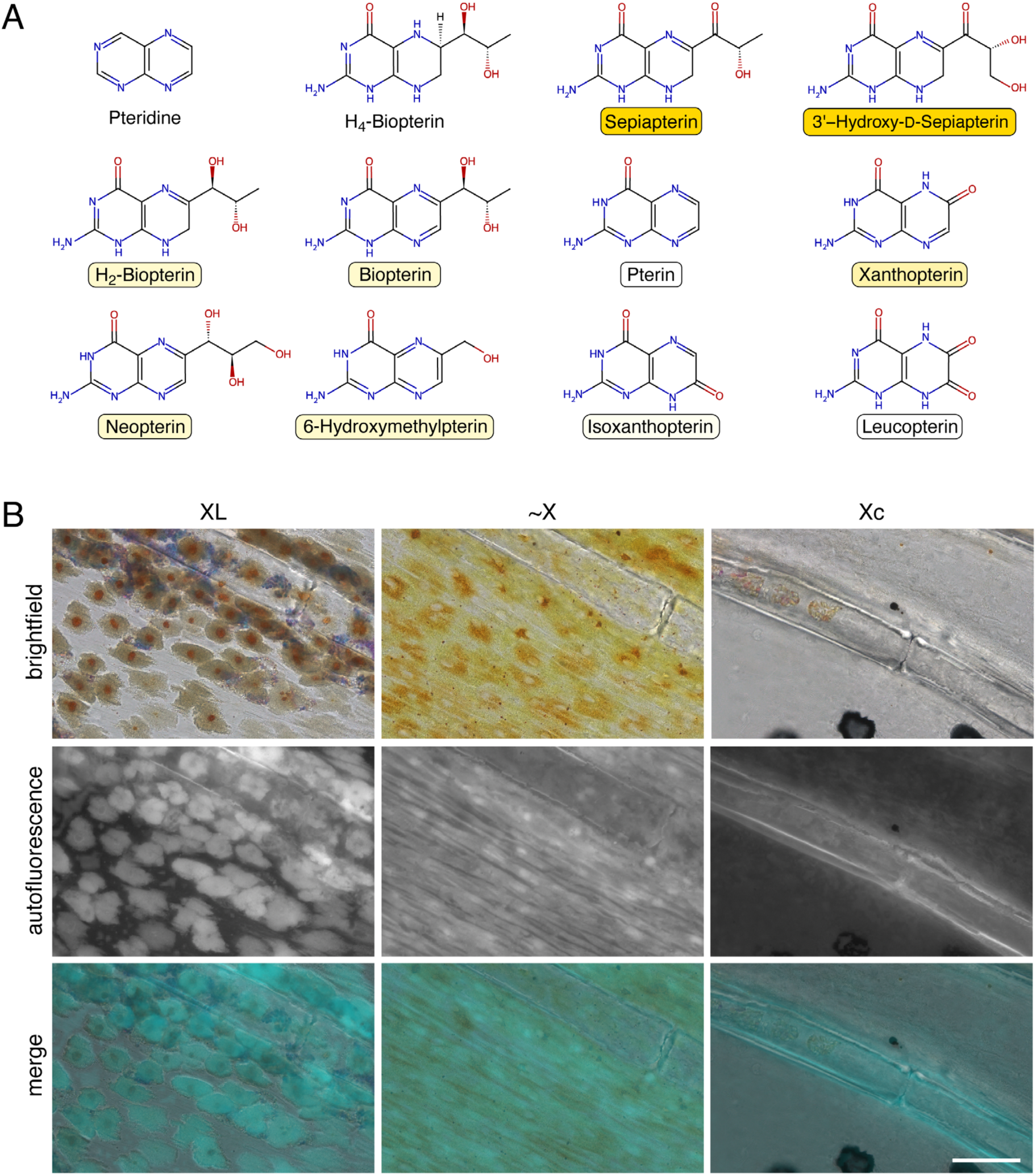
Pteridine-containing compounds and ammonia-induced autofluorescence. (*A*) The parent pteridine structure is shown at upper right, with tetrahydrobiopterin (H_4_-Biopterin), a product of sepiapterin reductase activity. Additional pteridines identified in this study are illustrated though many other such compounds have pigmentary functions, particularly in insects (6). Colors assignments derive from (6, 7) as well as descriptions of compounds purified commercially. (*B*) Many pteridines autofluoresce after treatment with dilute ammonia (8), as illustrated here for XL and ∼X in UV channel; a region of fin with cryptic xanthophores (Xc) had only minimal fluorescence.

**Fig. S8.**
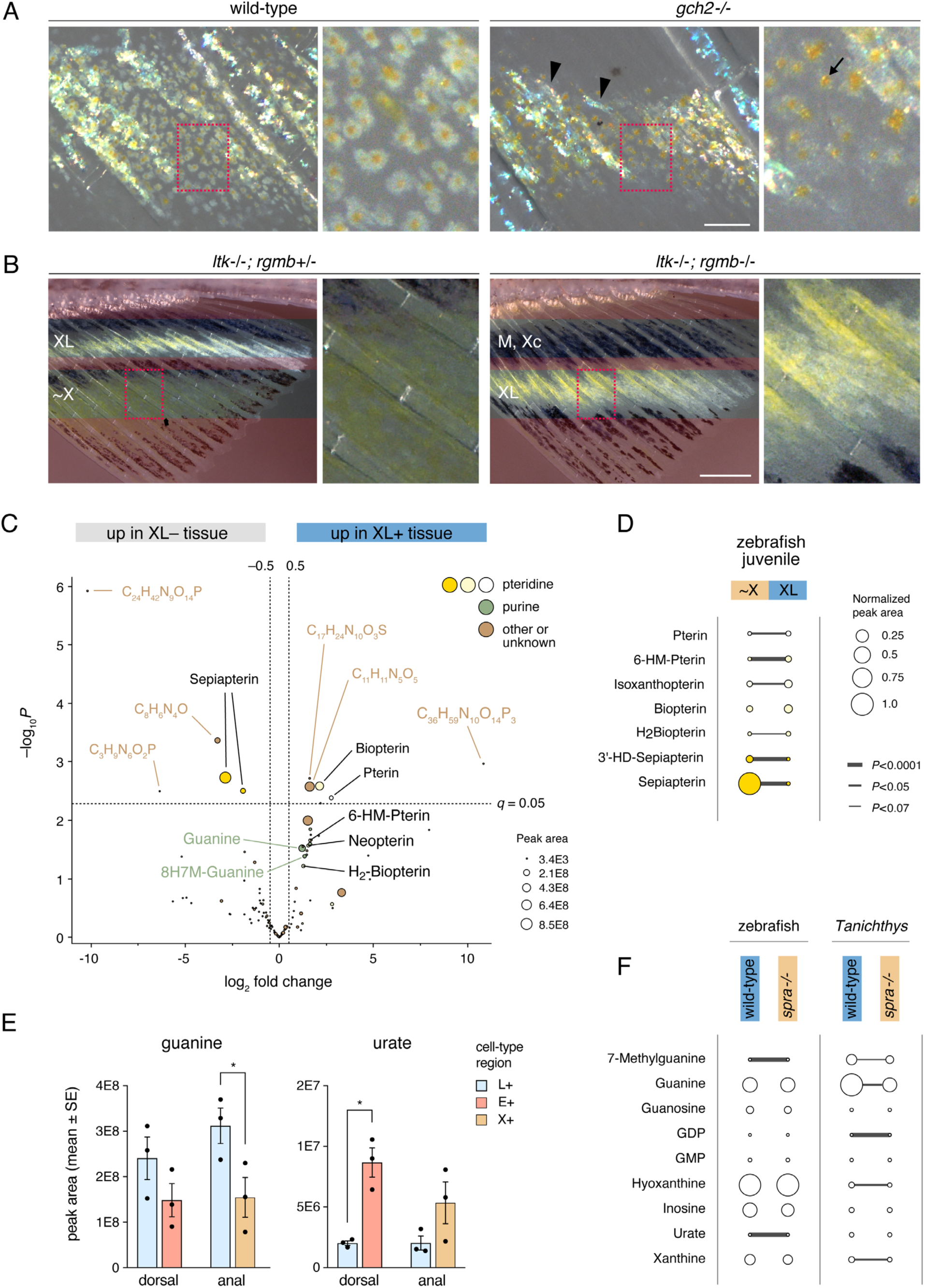
Pteridine pathway involvement in XL phenotype. (*A*) Zebrafish mosaic for CRISPR/Cas9 mutations in *gch2* had XL with reduced white material compared to wild-type and lacked the overall yellow appearance of XL in *spra-/-* mosaics. Neither iridophores (arrowheads) nor orange carotenoids of XL (e.g., arrow) were affected by *gch2* mutation. (*B*) Fins of zebrafish mutants for comparing tissue with or without XL (∼X, xanthophore-like cells; M, melanophores; Xc, cryptic xanthophores). Iridophores were missing due to *ltk* mutation whereas xantholeucophores were shifted distally due to a mutation in *rgmb* which was expressed more abundantly in XL than ∼X (log_2_FC=2.1, *q*=1E-15) (3, 9). Regions proximal and distal without red masks were dissected for metabolomic analyses. (*C*) Compounds differentially abundant between tissues lacking XL (∼X or melanophores with cryptic xanthophores) and tissues containing XL (without iridophores). Identifiable compounds indicated in black (pteridines), green (purines), or brown (other); compounds for which only chemical formulas could be determined shown in grey. (*D*) Pteridines found in metabolomic comparison of fin tissue from wild-type fish (14 SSL) containing ∼X (distal) or XL (proximal) showing average abundances (normalized to most abundant compound, sepiapterin); widths of connecting lines indicate *P*-values. (*E*) Targeted mass spectroscopy to evaluate guanine and urate abundance in *Tanichthys* dorsal or anal fin tissue with leucophores (L+), erythrophores (E+) or xanthophores (X+). * *P*<0.05 (*t*-test). (*F*) Metabolomic comparison of purines between wild-type and *spra-/-* mosaic fins of zebrafish and *Tanichthys*; key same as in *D*. Abbreviations: 6-HM-Pterin, 6-Hydroxymethylpterin; 8H7M-Guanine, 8-Hydroxy-7-methylguanine; GDP, Guanosine Diphosphate; GMP, Guanosine Monophosphate. (Scale bars: *A* 100 µm, *B* 500 µm)

## Materials and Methods

### Animals and Rearing Conditions

Fish were reared under standard conditions (∼28 °C; 14L:10D) with larvae initially fed marine rotifers derived from high-density cultures and enriched with Algamac (Reed Mariculture). Older larvae and adults were transitioned to live brine shrimp and flake food. Zebrafish stocks used were *chagall^vp36rc1^* (10), *duchamp^utr19e1^* (11, 12), *gch2^vc44^* (13), *gja5b^stl710^* (14), *ltk^j9s1^*, *rgmb^vp67re1^*, *Tg(aox5:palmEGFP)^wp.rt22^* (15) and *Tg(mitfa:Eos)* (generously provided by J. Lister). *Tanichthys albonubes* were obtained from The Wet Spot Tropical Fish (Portland, Oregon, USA). This study was performed in accordance with the recommendations in the Guide for the Care and Use of Laboratory Animals of the National Institutes of Health. Animals were handled according to approved institutional Animal Care and Use Committee (ACUC) protocol (#4170) of the University of Virginia. Euthanasia was accomplished by overdose of MS222 (Syndel) followed by physical maceration. For *Xenopus* oocyte assays, adult *X. laevis* were purchased from the Centre de Ressources Biologiques Xénopes (University of Rennes, France) and maintained at the AQUAB facilities of the Universitat Autònoma de Barcelona (UAB, Spain) as described (16). Oocytes were collected by surgical laparotomy from anesthetized females following a procedure approved by the Ethics Committee for Animal and Human Experimentation (CEEAH) from UAB and the Catalan Government (Direcció General de Polítiques Ambientals i Medi Natural; Project no. 10985).

### CRISPR/Cas9 Mutagenesis

CRISPR/Cas9 mutants or F0 mosaics were generated by injecting one-cell-stage embryos with approximately 1 nl of 5 µM gRNA:Cas9 RNP complex (IDT). For production of mutant lines, fish were screened for dorsal fin phenotypes at the juvenile stage and alleles recovered by incrossing and outcrossing. The *spra* mutants were lethal. Zebrafish *spra* genotyping was performed with: TAAATGTGAGGTTTTGATGC (forward primer), TAATCGATGCTGAACTCTC (reverse primer). *Tanichthys spra* genotyping was performed with: GTTTGCACTTTAGATCATGA (forward primer), TTAGTACATCCATCCACATG (reverse primer).

### Lineage tracing

Photoconversion was performed on *Tg(mitfa:Eos)* fish using a Zeiss LSM 800 laser scanning confocal microscope equipped with ZEN Blue software. Anal fin Eos+ xanthophore-like cells were photoconverted by 405 nm laser. Four to six cells per fish were converted in 7 fish. Following photoconversion, fish were maintained in tanks shaded from ambient light. Final imaging was performed first in fluorescence mode and then in brightfield mode. Although pigments exhibited autofluorescence in the same channels as Eos, treatment with epinephrine caused pigment granules to contract, allowing them to be distinguished from other Eos+ regions of cytosol.

### Imaging, Documentation and Analysis

Fish were anesthetized in MS222 prior to imaging. To contract pigment granules for imaging, fish were treated with 1 mg/ml epinephrine (E4642, Sigma-Aldrich) for 5 min. Bright-field and low-resolution fluorescent images of fish fins or bodies were acquired using Zeiss Axio Zoom stereomicroscope equipped with Zeiss Axiocam cameras, or Zeiss Axio Observer inverted microscope equipped with Yokogawa CSU-X1M5000 laser spinning disk and Hamamatsu camera. High-resolution fluorescence images were acquired using a Zeiss LSM880 inverted laser confocal microscope in Airyscan SR mode. Images were captured either as single frames or as tiled sets of larger areas that were then stitched computationally using ZEN Blue software. The same acquisition parameters were applied across matched sets of images (e.g., across genotypes and treatments) as were subsequent adjustments to color balance, display levels or both, applied to entire images as needed for visualization in Adobe Photoshop CS or ZEN Blue software.

### Pteridine fluorescence imaging

Fish were euthanized, and the anal fins were immersed in the imaging solution (2 drops of 5 M ammonia solution in 10 ml 1× PBS containing 0.1% β-mercaptoethanol, pH 10.0). Imaging was performed using the UV channel after anal fins were exposed to UV light for 10 s.

### Oil Red O staining

Anal fins were dissected and fixed in 4% paraformaldehyde for 2 h at room temperature. Samples were washed 3 times with 1× PBS for 5 min each, then incubated in 60% isopropanol for 20 min. Tissues were stained with 0.3% Oil Red O in 60% isopropanol overnight at room temperature. After staining, samples were washed with 60% isopropanol for 20 min, followed by three washes in 1× PBS for 5 min each, before imaging.

### Transmission Electron Microscopy

Fins were amputated and pre-fixed for 15–30 min with 4% formaldehyde and 2.5% glutaraldehyde in PBS, then 2 h in fresh 4% formaldehyde and 2.5% glutaraldehyde in 100 mM phosphate buffer at pH 7.4 at room temperature. Sample processing and imaging were as described (3).

### Metabolomic Analysis and High-Performance Liquid Chromatography

For zebrafish analysis, anal fin XL region, xanthophore region and cryptic xanthophore region were dissected from wild-type zebrafish, *ltk^j9s1^* mutants, *ltk^j9s1^*; *rgmb^vp67re1^* double mutants at 14 mm standard length (SL) stage. Three biological replicates were collected with each replicate containing fragments from 20 fish. For metabolomic analysis of zebrafish mutants, white regions of adult wild-type anal fins or the corresponding regions of *spra-/-* mosaic mutants were dissected. Three biological replicates were collected, with each replicate containing fragments from 6 fish. For *Tanichthys*, white regions of wild-type fish (fin tips of dorsal, anal, and pectoral fins) or the corresponding regions of *spra-/-* mosaic mutants were dissected. Three biological replicates were collected, with each replicate containing fragments from a single fish. Samples were collected on dry ice and ground into a powder using disposable pellet pestles under liquid nitrogen in pre-weighed tubes. Metabolites were extracted by adding 40 μL of ice-cold methanol:acetonitrile:water (2:2:1) solution per milligram of tissue, followed by two freeze-thaw cycles that consisted of immersion in liquid nitrogen for 1 minute followed by incubation at 25°C for 10 seconds, sonication for 5 minutes at 25°C, and vortexing for 30 seconds. Samples were incubated at -20°C for 1 h to precipitate protein and centrifuged at 20,000 x g for 10 minutes at 4°C.

Analysis of small molecules present in fin was performed on a Vanquish Flex UHPLC connected to an Orbitrap ID-X Tribrid mass spectrometer fitted with a H-ESI source (Thermo Fisher Scientific, Waltham, MASS). Metabolites were separated by using hydrophilic interaction liquid chromatography with a iHILIC column (100 x 2.1 mm, 5 µM, 200 Å (HILICON, Umeå, SWE)). Mobile phase A consisted of 95:5 water:acetonitrile, 20 mM ammonium formate, 0.1% ammonium hydroxide, 5 µM medronic acid and mobile phase B consisted of 95:5 acetonitrile:water were utilized with the following gradient: 0.0 – 1.0 min flow rate 0.250 uL/min and 90% B, 1.0 – 12.0 min 90 - 35% B, 12.0 – 12.5 min 35 – 25% B, 12.5 – 14.5 min 25% B, 14.5 – 15.0 min 25 – 90% B, 15.0 – 15.5 min 90% B, 15.5 – 16.5 min flow rate 0.250 – 0.400 mL/min and 90%B, 16.5 – 20.0 min 0.400 mL/min and 90%B, 20.0 – 20.5 min flow rate 0.400 – 0.250 mL/min and 90% B, 20.5 – 22.0 min 90%B. For MS acquisition source parameters were as follows: Sheath gas 50 (arb), Aux gas 10 (arb), Sweep Gas 1 (arb), Ion Transfer Tube 300 °C, Vaporizer Temp 200 °C) and the MS^1^ data was collected using the orbitrap detector (MS^1^ scan parameters: Resolution 120k, Scan Range 67 – 1000 m/z, RF Lens 60%, Normalized AGC Target 100%, Max injection time 200 ms, 1 Microscan, both polarities). A pool generated by combining equal amount from each sample was analyzed using MS^2^ Deep Scan method (Deep scan parameters: Orbitrap detector, MS^1^ resolution 60k, 1.5 m/z isolation window, stepped HCD collision energies (15, 30, and 45%), MS^2^ resolution 30k, Scan Mode auto, Max Injection Time 100 ms, Standard AGC target, 1 microscan). Retention times and MS/MS were collected on the following authentic standards; neopterin, isoxanthopterin, pterin, 6-biopterin, and 6-hydroxymethylpterin and compared to experimental samples to achieve level 1 identifications (17).

### Bulk RNA-Seq

Anal fin XL region and X region were dissected from ∼100 *Tg(aox5:palmEGFP)^wp.rt22^* of 1 month old (12∼14 mm SL). Fragments were dissociated to cell suspension by Liberase (Roche, Cat# 5401119001) at 30 °C for 30 min, followed by filtration through a 40 μm strainer. Isolated cells were sorted by fluorescence-activated cell sorting (FACS) using a Cytek Aurora CS (USA). GFP+ viable cells were collected, and RNA extracted using the RNAqueous™-Micro Total RNA Isolation Kit (Thermo Fisher, Cat. #AM1931). Sequencing libraries were prepared using the SMART-Seq mRNA LP Kit (Takara, Cat. #634768) and sequenced on an Illumina NextSeq 550 using the NextSeq 500/550 High Output Kit v2.5 (150 cycles). Reads were mapped and quantified using Kallisto (18) and differential expression analysis performed with DESeq2 (19). Pathway enrichment analyses were performed with https://www.webgestalt.org/ (20). RNA-seq data are available through GEO (accession: GSE313210).

### DNA Extraction

High molecular weight DNA was isolated following a protocol modified from (21). Tissues were incubated in 450 µL of Buffer B (0.4 M NaCl, 20 mM EDTA, 10 mM Tris-HCl pH 8.0) and 50 L of Buffer C (5% SDS, 2 mg/mL Proteinase K) at 55°C for 1.5–2 hours with intermittent vortexing, followed by overnight incubation at 55°C. Samples were treated with 2 µL RNase A (20 mg/mL) for 2–5 minutes. Protein precipitation was induced by adding 270 μL saturated NaCl (∼6 M) and pulse vortexing. Following centrifugation at 10,000 g for 30 minutes at 15 °C, the supernatant was transferred to fresh tubes containing 800 μL isopropanol. DNA was pelleted at 15,000 g for 25 minutes at 4°C, washed with 70% ethanol, air-dried, and eluted overnight at 4°C.

### Isolation and Molecular Cloning of *duchamp*

Fish were mutagenized with N-ethyl-N-nitrosourea using standard methods (22) and the duchamp allele identified as a dominant pigment phenotype in the F1 generation. After isolation and repeated crosses to the AB^wp^ inbred genetic background fish were crossed to the inbred mapping strain WIK and F1s in-crossed to generate a ∼2000 individual mapping panel segregating the *duchamp* phenotype. The mutant locus was placed on chromosome 5 by bulked segregant analysis and then fine-mapped to the vicinity of *aqp3a* on a subpanel of recombinant individuals using simple sequence repeats and other variants, with subsequent Sanger sequencing of exon sequences.

### Isolation and Mapping of *chagall* by Pool Sequencing

To identify the gene corresponding to *chagall*, mutants were crossed to the inbred mapping strain WIK and a single pair of F1s in-crossed to generate an F2 mapping family segregating the chagall mutant phenotype, from which mutant and wild-type pools containing DNA from 100 individuals of each genotype were prepared and sequenced. Sequences were concatenated and aligned to the *Danio rerio* reference genome (GRCz11) using Bowtie2 (23) with --sensitive-local parameters. The resulting BAM files were sorted, indexed, and processed with Samtools (24). Quality control of alignments was performed using Qualimap bamqc (25), assessing coverage and mapping quality. To reduce technical artifacts, we removed duplicate reads and filtered for high-quality alignments (mapping quality >= 20) using samtools markdup and view commands. Breadth of coverage was assessed by calculating the number of base pairs with a read depth of at least 5x. Variants were called using bcftools mpileup (26) with the -annotate AD,DP flag, followed by bcftools call. Variant calling was parallelized across genomic regions to maximize computational efficiency. The resulting BCF files were concatenated and subjected to initial quality control, checking for transition/transversion ratios, depth distribution, and missingness using vcftools. We applied a custom filtering script to retain high-confidence variants. Loci were filtered based on minimum mapping quality (MQ >= 30), base quality (QUAL >= 30), and minimum read depth (DP >= 2) per pool. For population genetic analyses, we computed F_st_, d_XY_, and pi in 1 kb sliding windows using pixy. Linkage disequilibrium (r^2^) was estimated using LDx.pl and averaged across 10 kb windows. We additionally implemented a custom “Dominant Score” algorithm adapted from (27). This score highlights regions exhibiting high heterozygosity in the mutant pool and homozygosity in the wild-type pool. The score was calculated in sliding windows of 100 SNPs with a step size of 20 SNPs using the formula:

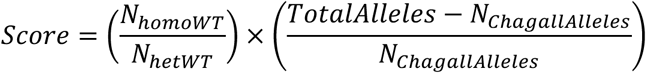

where counts were derived from genotype calls in the filtered VCF file. Significance thresholds were established via permutation tests (100 iterations), shuffling genotype values across chromosomes to define the 95th percentile cutoff.

Given that *chagall* is a dominant phenotype, we filtered for variants that were heterozygous in the mutant pool and homozygous reference in the wild-type pool. Known polymorphisms were excluded by filtering against the NHGRI-1 variant set and the Ensembl GVF database. Candidate SNPs and indels were intersected with gene annotations (Ensembl GRCz11 GTF) to identify exonic variants. Statistical association of candidate alleles with the phenotype was assessed using G-tests of independence, Chi-square goodness-of-fit tests, and Fisher’s exact tests. Final candidate variants were annotated for functional impact using VEP.

### Verification of *aqp3a* Mutant Alleles

Because *duchamp* and *chagall* mutants were phenotypically dominant and potentially exhibiting dominant negative or neomorphic activities we confirmed their correspondence to *aqp3a* by targeting exon sequence of *aqp3a* on the duchamp and chagall mutant backgrounds by CRISPR/Cas9 mutagenesis (IDT AltR Dr.Cas9.AQP3A.1.AA, target site CCAAACACTCGGATCCTTCT), with the expectation that induction of a premature termination codon would lead to loss of the dominant phenotypes. In each case mosaic, injected fish exhibited complete or partial rescue and recovered alleles genotyped to have both CRISPR-induced and ENU-induced mutations were phenotypically wild-type, as observed for other mutant alleles of *aqp3a* (2).

### Functional characterization Aqp3a mutant alleles

Water and glycerol permeabilities of wild-type and mutant Aqp3a were tested using the *X. laevis* oocyte expression system as previously described (28). Oocytes were injected with 15 ng of cRNA of each construct or not injected (controls). In some experiments, oocytes were co-injected with wild-type Aqp3a or Aqp3b (10 and 0.5 ng, respectively) plus 20 ng of the mutant forms. To determine the osmotic water permeability (*P*_f_), the oocytes were transferred at 48 h postinjection to 10-fold diluted MBS (88 mM NaCl, 1 mM KCl, 2.4 mM, NaHCO_3_, 0.82 mM MgSO_4_, 0.33 mM Ca(NO_3_)_2_, 0.41 mM CaCl_2_, 10 mM HEPES and 25 μg/ml gentamycin; 200 mOsm) at pH 8.5, and oocyte swelling was tracked by video microscopy. Glycerol permeability (*P*_gly_) was also determined volumetrically in isotonic Modified Barth’s Solution (MBS: where NaCl was replaced by 160 mM glycerol. The uptake of H_2_O_2_ was determined using the ROS-sensitive, cell-permeable fluorescent dye 5-(and-6)-chloromethyl-2′,7′-dichlorodihydrofluorescein diacetate, acetyl ester (CM-H_2_DCFDA; C6827, Life Technologies), following (29). The osmolarity of all solutions was measured with a vapor pressure osmometer (Vapro 5600, Wescor, USA), and adjusted to 200 mOsm with NaCl if necessary.

### Immunohistochemical localization of Aqp3a variants

Uninjected oocytes and oocytes expressing wild-type or mutant Aqp3a were processed for immunofluorescence microscopy(30). Section (8 μm) were blocked with PBST containing 5% normal goat serum and 0.1% BSA for 1 h at room temperature, and incubated with an anti-seabream (*Sparus aurata*) Aqp3a rabbit antibody (1:100 dilution in PBST) (31) overnight at 4°C in a humidified chamber. After washing three times with PBS, slides were incubated with sheep anti-rabbit IgG coupled with Cy3 (Merck, # C2306,) for 1 h at room temperature. Sections were washed with PBS and incubated (1:10000) with WGA Alexa Fluor® 647 conjugate (Life Technologies Corp., # W32466) for 10 min. The sections were mounted with fluoromount aqueous anti-fading medium (Merck, # F4680), and images were acquired at 20 × magnification with a Zeiss Axio Imager Z1/ApoTome fluorescence microscope.

### Whole Genome Sequencing and Assembly

We generated a high-quality genomic reference for *Tanichthys albonubes*, by processing PacBio HiFi reads at approximately 50x coverage with bamtools (32) to retain only high-quality sequences (rq >= 0.99) (Genbank BioProject ID PRJNA1393770, TaxID: 38762). We converted merged bam files to FASTQ format via pbindex and bam2fastq (available at https://github.com/PacificBiosciences/pbbam and https://github.com/jts/bam2fastq, respectively). Following quality visualization with NanoPlot (33), we *de novo* assembled the genome using hifiasm (v0.16.1) (34). The resulting assembly is highly contiguous: of 535 total scaffolds, 313 exceed 50 kbp, and 90% of the genome is contained within just 70 scaffolds (L90 = 70). We achieved an N50 of 18.40 Mbp and an N90 of 3.96 Mbp; notably, the largest contig spans 39.57 Mbp, approaching the chromosome scale of the related model teleost *Danio rerio* (∼50 Mbp, 2n=50). We assessed the completeness of our *de novo* assembled genome we used BUSCO (v5.8.3) (35) against the actinopterygii_odb10 lineage. The *T. albonubes* genome shows 99.0% completion, with 94.8% single-copy, 4.2% duplicated, 0.5% fragmented, and 0.5% of missing genes. Finally, we generated an annotation for *T. albonubes* by lifting *D. rerio* annotations to the masked *T. albonubes* assembly with Liftoff (36) (parameters: -polish -flank 0.25 -d 2.5). We evaluated synteny by generating whole-genome alignments with nucmer (MUMmer4)(37) filtering coordinates via show-coords (-B -r -I 80 -L 1000), and identifying bundles with bundlelinks (maximum gap 10 kbp, minimum size 1 kbp) for visualization in R using ggplot2 and GenomicRanges.

### Statistical Analyses

Analyses of quantitative data were performed in JMP Pro 18 (SAS Institute, Cary NC) or for metabolomic data using the limma package in R (38) for improved variance stabilization and cross-feature variance estimation using a multifactorial model to control simultaneously for genotype, anatomical position and replicate in estimating effects of cell-type composition across tissue samples.

